# Coping strategies dynamics and resilience profiles after early life stress revealed by behavioral sequencing

**DOI:** 10.1101/2025.09.01.673507

**Authors:** Jeniffer Sanguino-Gómez, Umut Güçlü, Harm J. Krugers, Antonio Lozano

## Abstract

Animal models can provide valuable insights into the mechanisms underlying stress-related disorders. Yet, significant translational challenges persist, as laboratory behavioral assays are often reductionistic, with limited attention to ethologically relevant behavioral diversity. Recent advances in high-throughput pose-estimation tools and computational ethology methods are addressing this limitation by enhancing the resolution and validity of behavioral phenotyping. In this context, it is known that early life stress (ELS) reshapes how animals handle subsequent threats later in life, but the fine-scale dynamics and ethological details of this shift remain elusive. To overcome this, we combined a deep-learning pose-estimation pipeline (DeepLabCut) with a supervised freezing classifier (SimBA) and an unsupervised behavioral motifs identification platform (keypoint MoSeq) to study in detail the diversity and dynamics of behavior in an auditory fear-conditioning (FC) paradigm in two independent cohorts of adult male mice that were exposed to ELS through the limited bedding and nesting (LBN) paradigm. We first validated the blunted freezing response after ELS in a supervised manner using SimBA. Next, keypoint MoSeq segmented the same pose-estimation data into ethologically meaningful motifs over time. When compared to control animals, ELS offspring showed an altered FC response, reduced behavioral entropy and limited diversity in their behavioral repertoire. Such response was characterized by longer active-behavior bouts and more recurrent transitions between states, indicating a more stereotyped and predictable response. Multidimensional scaling of time-binned behavioral vectors and distance metrics identified a resilient subpopulation within the ELS group that displayed a control-like behavioral profile, characterized by a steeper increase in freezing behavior during the FC task and a more diverse behavioral repertoire with reduced recurrence of stereotyped actions, less frequent and shorter active bouts and prolonged passive responses. Overall, our findings suggest that ELS shifts the balance between passive and active coping strategies and that resilience is marked by a less stereotypical yet more diverse and flexible behavioral response to a subsequent stressful demand. Finally, we further validated the unsupervised behavioral motifs with a predictive model that identified distinctive kinematic features of these responses, which could be used to build new behavioral classifiers that can be applied in other behavioral paradigms. These results demonstrate the potential of computational ethology to dissect complex behavioral patterns and improve our understanding of individual stress responses. By combining supervised and unsupervised behavioral analysis tools, we can deepen our understanding of the latent structure of stress behavior and identify objective markers of vulnerability and resilience.

## 1 Introduction

Exposure to stressful situations triggers a behavioral response, classically defined by Cannon^1^ as the *fight or flight* response-characterized by sympathetic nervous system activation and corticosteroid release into the bloodstream-preparing the body to confront or escape danger. As survival depends mainly on the ability to successfully cope with the threat^2^, coping strategies are therefore essential in determining the outcomes of aversive life events^2^.

While the *fight or flight* response represents the ingrained survival instinct, its generalizability across individuals and contexts is limited. Subsequent research has expanded this dichotomous response to include additional coping strategies, such as freeze or fawn^3,4,5,6^. Coping refers to these response patterns that occur in reaction to a challenging environment^7^ and can be broadly categorized into active (*fight or flight*) or passive (decreased responsiveness to the environment, immobility) behavioral strategies, based on the presence or absence of attempts to act upon the stressor^5,2^.

Effective coping with stressful events requires appropriate cognitive processing to evaluate the situation and promote behavioral adaptation^8,9^. Coping can be influenced by various individual factors, such as genotype, sex, or (early-life) experiences^10,11,12,13,14,15^. In particular, the impact of stressful life events on physical and psychological adaptive responses varies significantly among individuals^16,17,18^.

Some of the (mal)adaptive responses to stressors may serve as predictors of stress-related disorders, including anxiety and post-traumatic stress disorder (PTSD)^19,20,21,22^. Reliably measuring the full spectrum of behavioral responses and coping strategies in depth may help understanding resilience and vulnerability.

In response to the challenges in behavioral profiling, recent advancements in deep learning pose estimation tools, such as DeepLabCut^23^, have revolutionized the field of behavioral neuroscience. These pose estimation tools enable precise tracking of user-specified points, allowing for more complex and detailed behavior analysis compared to traditional tracking methods or manual scoring. The resulting post-estimation datasets allow researchers to extract and characterize behavioral traits through supervised methods (among others:^24,25,26,27,28,29^) or to explore data patterns without external information using unsupervised approaches (among others:^30,31,32,33,27,34,35,36,37^). Here, we examined whether and how a previous history of stress-exposure during early development, early-life stress (ELS), shapes later behavioral coping strategies in response to a subsequent threat and whether it alters behavioral signatures associated with susceptibility and resilience. To achieve this, mice underwent ELS using a standardized model involving the restriction of bedding and nesting (limiting bedding and nesting model^38^). In adulthood, these animals were subjected to classical auditory fear conditioning (FC)^39,40,41,42^, where a conditioned stimulus is paired with an aversive unconditioned stimulus. This widely used laboratory model of fear learning allowed us to study coping strategies in the face of a threat. In this task, rodents instinctively freeze in response to the footshock, making freezing a suitable supervised category^43,44,45^. Preliminary data demonstrated the potential of a random forest classifier, SimBA^24^, to automatically score freezing using DeepLabCut pose estimation data (implemented in^46,47^). We validated this classifier and further applied an unsupervised pipeline to explore the full behavioral repertoire^48^ of these animals without pre-training, offering a data-driven approach that may reveal previously unrecognized, unbiased coping strategies. While existing literature suggests that ELS animals exhibit reduced freezing behavior during FC^49,46,47,50^, we hypothesized that a previous history of ELS also alters the broader structure of behavior during threat exposure. These differences may be reflected in the duration, frequency, transitions and/or entropy of behaviors. Our approach therefore aimed to capture a richer, dynamic spectrum of the behavioral responses to stress-going beyond measuring freezing behavior but also to include subtle emergent behavioral patterns that traditional analysis methods often overlook. Finally, this pipeline further enabled a detailed characterization of individual differences in behavioral responses, providing a powerful tool to distinguish distinct signatures of susceptibility and resilience shaped by early adversity.

## 2 Material and methods

### 2.1 Animals and breeding

Data from previously published studies^46,47^ was used, and as such, they were conducted in a transgenic mouse model expressing a fluorescent reporter gene under the promotor of the immediate early gene Arc (Arc::dVenus mice, kindly provided by prof. dr. Steven Kushner, Erasmus University Rotterdam -^51,52,53,54^).

All mice were housed under standard conditions (20–22 °C, 40–60% humidity) with a 12-hour light/dark cycle (lights on from 8 a.m. to 8 p.m.) and continuous background radio noise. Standard chow and water were available *ad libitum*. All experiments were carried out during the light phase. All experiments were approved by the Animal Welfare Committee of the University of Amsterdam, in accordance with EU directive 2010/63/EU.

To ensure a standardized perinatal environment, mice were bred in-house. Six to eight-week-old female C57BL/6J mice were purchased from Envigo Laboratories (Venray, Netherlands) and acclimatized during a two-week period before breeding started. For mating, two females and one Arc::dVenus homozygote male mouse were co-housed in a conventional type II cage with standard bedding, nesting material, and paper enrichment for a period of one to ten days to facilitate mating^51,53^. Wild-type females were deliberately chosen for breeding to mitigate potential line-dependent variations in maternal care.

Following mating, the female mice continued to be housed together for an additional week with standard bedding, a reduced amount of paper cage enrichment, and a 5 x 5 cm square of cotton nesting material (Tecnilab-BMI, Someren, The Netherlands). After this paired-housing period, pregnant primiparous females were individually housed in a conventional type II cage, covered with a filter top, containing normal bedding, one nestlet, and a modest amount of retained old bedding material, without paper cage enrichment. The animals were transferred to a different room starting 18 days after the initiation of breeding and were monitored each day for the birth of the pups. Dams were checked for the birth of pups each morning before 09:00 a.m. When a litter was born, the previous day was designated as PND 0.

Total numbers of animals per experiment are summarized in **Table 1**. One animal was excluded from the analysis due to its failure to recapitulate ground truth freezing, as identified by the supervised classifier. Visual inspection indicated that this discrepancy may have resulted from inaccurate pose estimation tracking.

**Table 1.** Number of animals per experiment and group.

| Experiment | Control | ELS | Total |
| --- | --- | --- | --- |
| Sanguino-Gómez and Krugers (2024) | 25 | 25 | 50 |
| Sanguino-Gómez et al. (2024) | 16 | 16 | 32 |
| **Total** | 41 | 41 | 82 |

### 2.2 Early life stress paradigm

The early life stress paradigm consisted of housing mice with limiting bedding and nesting (LBN) material between PND 2 to PND 9 as described previously^55,56,38^. At PND 2, litters were culled to 6 pups per litter. Litters with fewer than five pups or consisting of only one sex were excluded from the analysis. Dams and their litters were randomly allocated to either the ELS or control condition until PND 9, after which all mice were treated equally (**Fig. 1A**).

**Figure 1.**
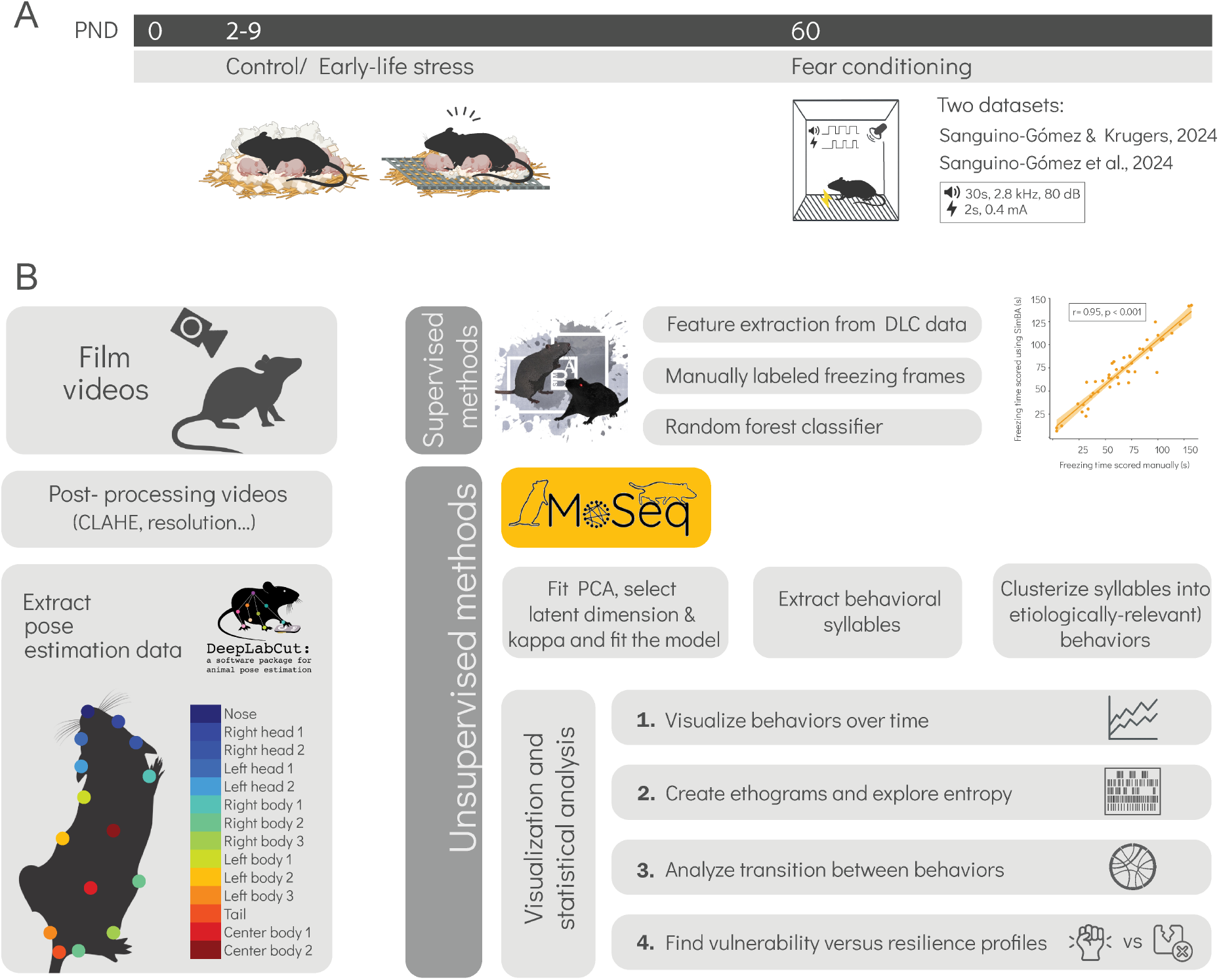
Experimental design and analysis workflow. **A**. Schematic overview of the auditory fear conditioning experiment. Male mice were subjected to either standard conditions or ELS from PND 2–9. At PND 60, mice underwent training in a mild auditory fear conditioning task (Tone: 30 s, 2.8 kHz, 80 dB, footshock 2 s, 0.4 mA) in two independent cohorts ^46,47^. Partially created with BioRender.com. **B**. Analysis workflow. Raw videos were preprocessed (resolution adjustment and CLAHE) and pose-estimated with DeepLabCut using a 14-point mouse model. In the supervised pipeline using Simple behavioral analysis (SimBA), kinematic features (speed, angles, inter-body parts distances, etc.) were extracted from pose estimation data and aligned to manually labeled freezing frames. A random-forest classifier trained on these features achieved R = 0.95 against manual annotations, enabling accurate automated freezing prediction. In the unsupervised pipeline with Motion sequencing (keypoint MoSeq), pose estimation data was reduced via principal component analysis, the optimal latent dimensionality was chosen by *κ*-tuning and an autoregressive hidden Markov model segmented behavior into syllables. Those syllables were then clustered into ethologically relevant motifs. Downstream visualization and statistics include (1) time-series plots, (2) ethograms and entropy metrics, (3) transition network analyses, and (4) identification of vulnerability versus resilience profiles.

The control group was provided with a standard amount of sawdust ( ≈ 100g) and a standard 5 x 5 cm square of cotton material (Tecnilab-BMI, Someren, The Netherlands). In contrast, for the ELS group, dams were supplied with a reduced amount of sawdust ( ≈ 33g, one-third of the standard amount) and half of the standard square of cotton material (2.5 x 5 cm). Additionally, a stainless-steel grid (22 cm x 16.5 cm, mesh size 5 mm x 5 mm, in-house made) was placed 1 cm above the cage floor to prevent the use of sawdust for nesting. Filter tops were maintained for both conditions. Both groups were left undisturbed until PND 9, when mice were weighed and returned to standard housing conditions, where they remained until weaning at PND 21. At this timepoint, ear clips were obtained for identification purposes and to determine their genotype.

### 2.3 Fear conditioning

Two months old (± 2 weeks) male mice underwent testing in an auditory fear conditioning paradigm (**Fig. 1A**). To prevent potential bias in subsequent testing order and the transmission of fear, the mice were individually housed in isolated, sound-proof cabinets one week before fear conditioning. All behavioral procedures were conducted in the morning, specifically between 08:00 a.m. and 10:00 a.m., when basal plasma corticosterone levels are typically low.

The fear conditioning protocol was run using a computer with Ethovision software (version 14.0, Noldus, The Netherlands) and mouse behavior was recorded using a Basler acA1300-30gm GigE camera (Ahrensburg, Germany) connected to the computer. On the training day, mice were introduced to the conditioning chamber (Context A: 17 cm x 17 cm x 25 cm, Ugo Vasile, with a steel grid floor connected to a shock generator) for 180 seconds, during which they were exposed to white noise (1.2 kHz, 50 dB) to freely explore the context. Following the exploration period, a tone (Conditioned Stimulus, CS: 30 s, 2.8 kHz, 82 dB) was presented, paired with a foot shock (Unconditioned Stimulus, US) during the last two seconds of the tone (2s, 0.4 mA). This tone-shock pairing was repeated three times, with each repetition separated by an intertrial interval of 60 seconds. Sixty seconds after the last foot shock, the mice were returned to their home cage. After every trial, the conditioning chamber was cleaned with 96% ethanol to eliminate residual odors from previous trials that could influence subsequent animal performance.

### 2.4 Data preprocessing

Videos of mouse behavior were adjusted to a resolution of 800 x 600 pixels and a frame rate of 25 frames per second (FPS). As a method for enhancing the local contrast of the images, we pre-processed the videos using Contrast Limited Adaptive Histogram Equalization (CLAHE -^57,58^. **Fig. 1B**). CLAHE is a non-linear image enhancement technique that operates by dividing an image into small, overlapping regions and then applying histogram equalization independently to each of these regions, ensuring that the contrast enhancement is performed locally. To prevent excessive amplification of noise during the equalization process, CLAHE incorporates a contrast limiting mechanism, restricting the enhancement within a predefined range. This method aims to enhance local details without compromising overall image quality.

### 2.5 Pose estimation and tracking

To build our pose estimation model, we used DeepLabCut (version 2.1.8.2 -^23^ - **Fig. 1B**). A total of 2000 frames were extracted from an independent sample of 100 videos, with 90% of labeled frames allocated to the training set and the remaining 10% included in the test set for model evaluation. The model was trained for 100000 iterations, saving a snapshot every 1000 iterations, using ResNet-50 with a batch size of 32 frames. Performance was evaluated at each snapshot by calculating loss, a measure of the error between predicted and true values (user labels) every 1000 iterations. Snapshot at 100000 iterations showed an average error of 2.76 pixels on training frames and 4.64 pixels on test frames after filtering for points with a likelihood cutoff threshold greater than 0.6. Visual inspection of labeled held-out videos confirmed the model’s performance. Data re-labeling was performed on low likelihood frames where the model initially failed to correctly track the body part.

The model tracked 14 body points, including the nose, left and right ears, three points per each body side, two points along the spine and the tail base (**Fig. 1B**). These points provided adequate coverage of the animal regardless of partial body occlusion or orientation relative to the camera.

The DeepLabCut model will be available as supplementary data in the code and data availability statement.

### 2.6 Supervised training of freezing classifier

Simple Behavior Analysis (SimBA)^24^ was used to generate a freezing behavior classifier from pose estimation data generated through DeepLabCut from a subset of 33 videos (**Fig. 1B**). First, 216 features were extracted, including body part locations (X, Y coordinates), movement metrics, inter-body part distances, confidence probabilities, and deviation-based measures. These features captured positional data, frame-to-frame displacement, Euclidean distances between body parts, and probability scores from pose estimation. We generated a dataset using manually annotated videos were to establish ground truth labels for each frame. A total of 182918 frames from 33 videos from an independent sample were used to train the model for freezing behavior detection. We used a 80/20 training/validation split. A random forest model was trained using SimBA, with 1000 estimators, a minimum node size of 20, an RF criterion of gini, a maximum feature selection of sqrt, and SMOTEEN oversampling to address class imbalance. We established a minimum bout size of 250 milliseconds to consider the behavior to be labeled as freezing.

Model performance was assessed by calculating precision, recall, and F1 scores. The classification report showed a precision of 0.937, recall of 0.865, and F1 score of 0.900 for freezing behavior, based on 8709 frames. For non-freezing behavior, the model achieved a precision of 0.957, recall of 0.981, and F1 score of 0.969 across 26521 frames.

After training and evaluating the model, we used it to predict the freezing behavior of an independent manually annotated set of videos, with a total of 211500 frames (**Fig. 1A**).

The SimBA model will be available as supplementary data in the code and data availability statement.

### 2.7 Unsupervised behavioral motifs discovery with keypoint MoSeq

To identify behavioral syllables in an unsupervised manner, we trained an autoregressive hidden Markov model (AR-HMM) on our DeepLabCut dataset using keypoint MoSeq^37^. Before fitting, keypoint MoSeq translated and rotated body-part coordinates into a common egocentric reference frame and compressed them into a six-dimensional latent space. It then fitted a vector-AR-HMM with a temporal stickiness hyper-parameter *κ*, empirically tuned to 1 × 10^6^ to produce syllable lengths of roughly 400 ms. Training occurred in two stages. First, we ran 50 Gibbs-sampling iterations on an AR-only model to initialize emission parameters and refine *κ*. Next, we performed 450 full AR-HMM iterations to jointly estimate transition probabilities, syllable centroids, heading vectors, noise covariances and continuous latent trajectories. The final model described 96 syllables, which after filtering by occurrence to capture 99.5% of labeled frames, 38 syllables remained for downstream analysis.

### 2.8 Grouping behaviorally relevant syllables

To interpret the behavioral relevance of the keypoint MoSeq syllables, we first visually inspected each syllable’s average pose trajectory over representative video segments. We then assigned qualitative behavioral labels and grouped syllables into seven behavioral clusters, ranging from more passive to more active behaviors: freezing, sniffing, grooming, turn, locomotion, climbing and jump. Behavior definitions are listed in **Supplementary Table 1**. Additionally, we identified syllables representing mixtures of multiple behaviors (e.g., transitions) and those with noisy tracking. These were grouped as “unassigned” frames.

During the visual inspection, we observed that climbing behavior could not be reliably captured by keypoint MoSeq’s predictions in its default configuration, likely due to the inherent limitations of two-dimensional top-down video in representing vertical movements. To isolate climbing events, we generated a binary mask separating the floor and walls of the cage based on manually annotated corner points from the median background frame of each video. For each frame, we then computed the convex hull of the animal’s DeepLabCut keypoints and quantified the fraction of its area overlapping with the floor (**Supplementary Figure 2**). Frames with a floor-overlap ratio above 0.8 were considered as potential climbing frames. To avoid false positives, we grouped potential climbing frames into bouts longer than 425 ms (17 frames) and labeled them as climbing. Shorter bouts were reassigned to the dominant non-climbing label within the sequence.

### 2.9 Frequency-based metrics and bout duration analysis

Behavioral data with clustered assignments per 250 milliseconds were used for further analysis. For each animal, we calculated the relative frequency of clusters and derived frequency-based measures including Simpson’s diversity index, Shannon entropy and evenness, as described below:

#### Simpson diversity score

The Simpson diversity score (*D*) (**Equation 1**) was computed to quantify the overall behavioral diversity exhibited by each animal. This measure represents the probability that two randomly selected time bins from the same animal would belong to different behavioral clusters. For each animal, the relative frequency (*p*_*i*_) of each observed behavioral cluster was calculated by dividing the number of time bins assigned to a specific cluster by the total number of observed bins. The diversity score was then calculated using the formula:

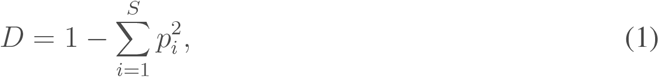

where *S* denotes the total number of behavioral clusters observed for each animal. Scores approaching 0 indicate low behavioral diversity, characterized by behavior concentrated in fewer clusters, while scores approaching 1 indicate high behavioral diversity, reflecting engagement across many different behaviors.

#### Shannon entropy

The Shannon entropy (*H*) (**Equation 2**) was computed to quantify the uncertainty or information content of the behavioral repertoire exhibited by each animal. This measure reflects how evenly distributed the behaviors are, with higher entropy values indicating a more diverse and balanced distribution, and lower values suggesting dominance by a few behaviors. For each animal, the relative frequency (*p*_*i*_) of each behavioral cluster was determined by dividing the number of time bins assigned to that cluster by the total number of observed bins. The entropy was then calculated using the formula:

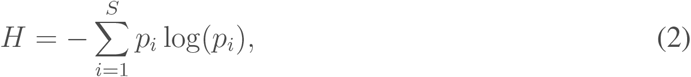

where *S* denotes the total number of behavioral clusters observed for that animal. When all clusters are equally likely, the entropy is maximized; when one or a few clusters dominate, the entropy is lower.

#### Evenness

Evenness (**Equation 3**) quantifies how uniformly the behavioral clusters are represented relative to the maximum possible diversity. It is computed as the ratio of the observed Shannon entropy to the maximum entropy, which is log(*S*) (the entropy value when all *S* clusters are equally represented). The evenness value (*E*) is given by:

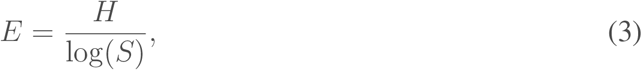

Values of *E* range from 0 to 1, with values near 1 indicating that behaviors are very evenly distributed among clusters, and values closer to 0 indicating an uneven distribution dominated by one or a few behaviors.

#### Cumulative usage index

The Cumulative Usage Index (CUI) (**Equations 4** and **5**) was computed to quantify how quickly an animal’s behavioral repertoire is covered by its most frequently used clusters. For each animal, the relative usage *p*_*i*_ of each behavioral cluster was determined by dividing the number of time bins assigned to that cluster by the total number of observed bins. These clusters were then ordered from highest to lowest relative usage. Let *p*_1_, *p*_2_, …, *p*_*S*_ represent the sorted relative usage values for the *S* behavioral clusters observed for each animal. The cumulative sums *c*_*i*_ of these usage values were computed as:

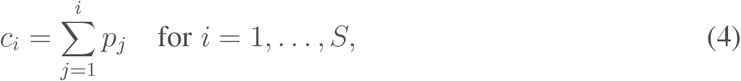

and the CUI was then calculated by taking the average of these cumulative sums:

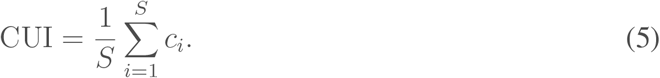

Values of CUI approaching the lower end indicate that a few clusters dominate the animal’s behavior, whereas higher values suggest a more even distribution of behavioral usage across the repertoire.

#### Bout duration

Bout duration was analyzed by identifying consecutive time bins with constant cluster assignments. Overall and cluster-specific mean bout durations were computed per animal.

### 2.10 Transition-based metrics

For each animal, the transition-based metrics were computed directly from the raw sequence of cluster assignments obtained from the behavioral sequencing recorded every 250 milliseconds.

#### Lempel–Ziv complexity

For an animal with a sequence of cluster assignments

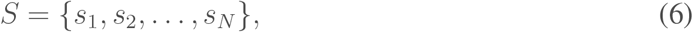

the Lempel–Ziv complexity (**Equations 6** and **7**) quantifies the diversity of transition patterns in the behavioral sequence. The algorithm scans *S* from left to right and counts the number of new substrings encountered. Starting with a counter *C* = 1 (to account for the first state), the sequence is parsed to identify the shortest substring that has not been previously observed. Each time a new substring is found, *C* is incremented. Although there is no simple closed-form formula, the computed measure is given by:

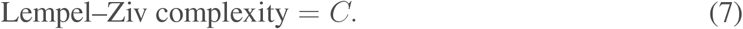

Higher values indicate a richer repertoire of transitions.

#### Recurrence rate

The recurrence rate (**Equations 8** and **9**) quantifies how frequently the same behavioral state recurs over time. A recurrence plot *R* is first constructed using the rule:

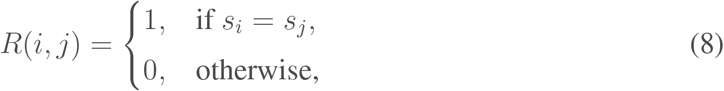

for *i, j* = 1, …, *N*. The recurrence rate is then calculated as the proportion of points in the plot that indicate a recurrence:

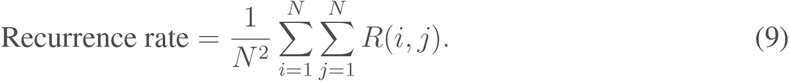

#### Determinism

Determinism (**Equation 10**) measures the predictability of the behavioral sequence by quantifying the fraction of recurrent points that form diagonal line structures in the recurrence plot. These diagonal lines represent sequences of consecutive repeated states. Let {*L* = *l*_1_, *l*_2_, …, *l*_*k*_} be the set of lengths of diagonal lines (with a minimum length threshold, typically 2). Determinism is defined as:

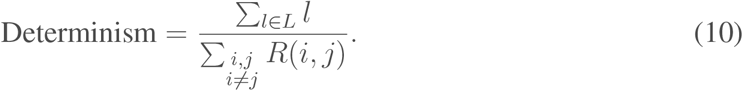

Higher determinism values suggest that the sequence has more predictable, structured transitions, whereas lower values indicate a more random sequence.

#### Markov entropy

Markov entropy (**Equation 11**) quantifies the unpredictability of behavioral transitions over time. For each animal, the metric was computed directly from the sequence of cluster assignments derived from behavioral recordings sampled every 250 milliseconds. Each sequence was treated as a first-order Markov process, in which the transition probability from one behavioral cluster to the next depends only on the current state.

To compute this measure, transition counts between all pairs of clusters were first tabulated. A small constant was added to each transition count to ensure numerical stability (Laplace smoothing), after which the counts were normalized to obtain a transition probability matrix *P*, where *P* (*i, j*) reflects the probability of transitioning from cluster *i* to cluster *j*. In parallel, the stationary distribution *π* was estimated as the empirical proportion of time each cluster was occupied.

The Markov entropy was then calculated as the expected conditional entropy of future states given the current one, using the formula:

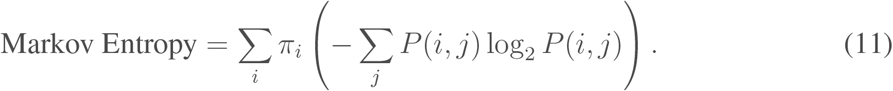

This value represents the average uncertainty in predicting the next behavioral cluster. Higher entropy values indicate more diverse and less predictable sequences, whereas lower values reflect greater regularity or repetition in the behavioral transitions.

### 2.11 Behavioral dynamics score

For each animal we generated cluster sequences by binning the behavioral data into 30-second intervals (750 frames per bin) and computing the percentage of time spent in each behavior within each bin. These binned percentages were then flattened into a single feature vector representing the relative frequency of each behavioral cluster over time (“behavioral profile”, **Fig. 2**).

**Figure 2.**
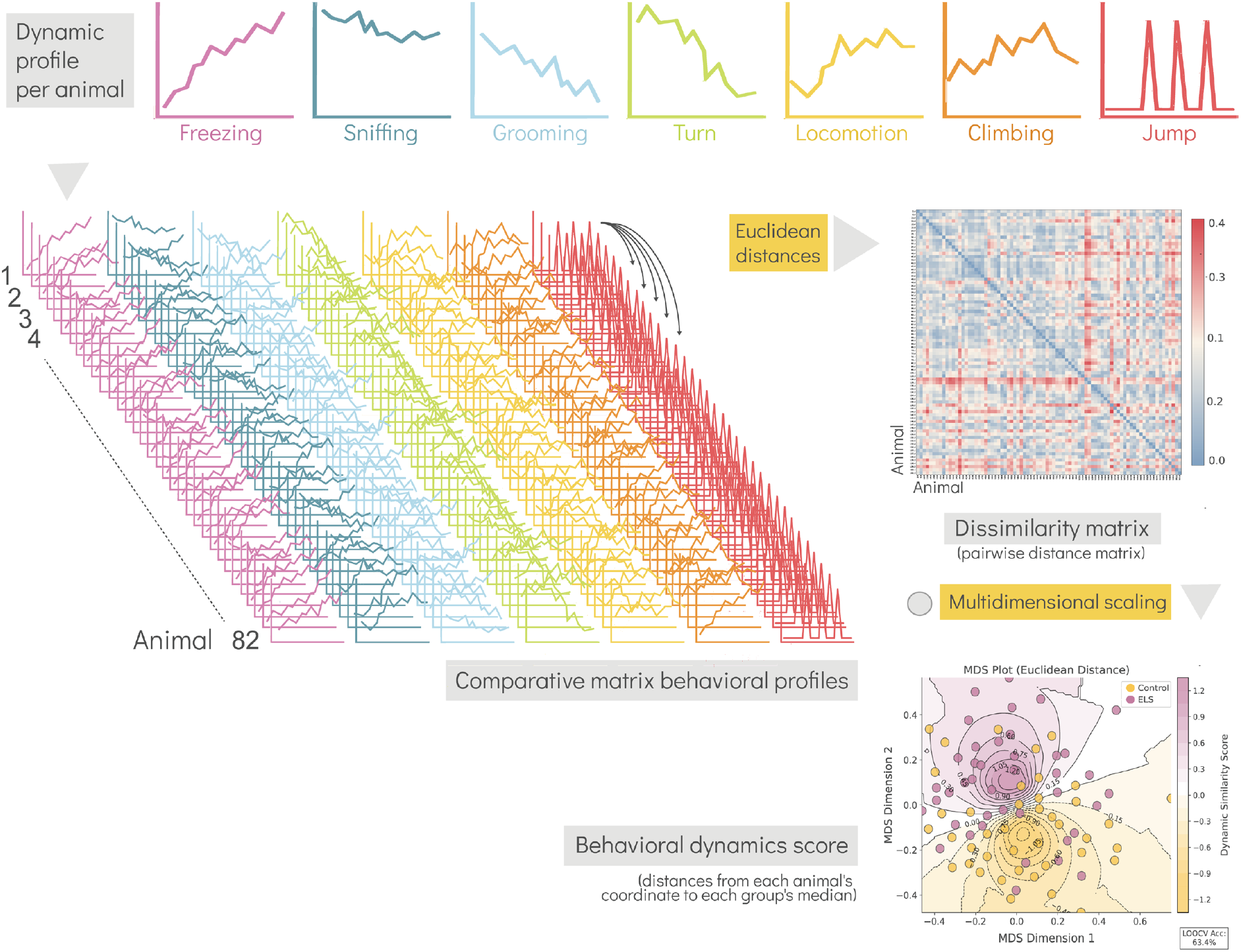
Behavioral-dynamics profiling and similarity mapping. We first compute a dynamic behavioral profile for each mouse by binning the FC session into 30-s intervals and calculating the percentage of time spent in each motif cluster (freezing, sniffing, grooming, turn, locomotion, climbing, jump). Next, we flatten each profile into a feature vector and stack them into a comparative matrix to align every animal against all others. From that matrix, we compute a dissimilarity matrix (pairwise Euclidean distances) to quantify how each animal’s profile differs from the rest. We then apply MDS to project the dissimilarity matrix into a coordinate space, positioning each mouse in two dimensions so that Euclidean distances in this space mirror the original profile dissimilarities. We calculate each group’s median coordinate in MDS space and, using these, derive a behavioral dynamics score for each mouse as the log ratio of its Euclidean distance to the ELS median versus its distance to the control median. Positive scores indicate that an animal’s profile lies closer to the ELS group (vulnerability), while negative scores indicate closer alignment to the control group (resilience). This is represented in the behavioral similarity landscape (MDS embedding), placing each animal along a vulnerability–resilience axis.

Pairwise distance matrices were computed using Euclidean distance on these feature vectors. For validation, we also computed distance matrices using Cosine, Jensen–Shannon, Manhattan and Correlation metrics (**Supplementary Figure 3**) based on the relative frequencies of all observed clusters.

To visualize the behavioral space, we applied multidimensional scaling (MDS) (**Equation 12**) to the distance matrix. MDS was computed by finding a two-dimensional configuration **Y** = {**y**_1_, …, **y**_*n*_} that minimizes the following stress function:

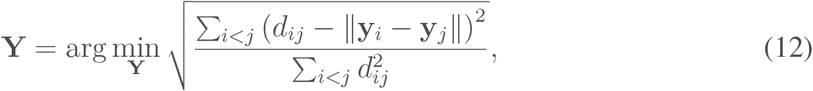

where *d*_*ij*_ are the normalized Euclidean distances computed from the feature vectors. This produced a two-dimensional representation of the animals’ behavioral profiles (**Equation 12**).

Within the resulting MDS space, we then computed the behavioral dynamics score (**Equation 13**) as follows (inspired by^36^):

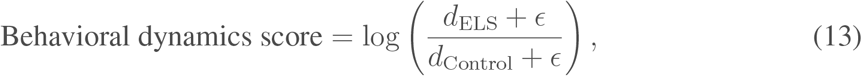

where *d*_ELS_ and *d*_Control_ are the Euclidean distances from each animal’s coordinate to the median coordinate of the ELS and Control groups, respectively, and *ϵ* is a small constant to avoid division by zero. A schematic overview of the entire workflow is presented in **Fig. 2**.

Animals were classified as resilient or vulnerable using a cutoff of zero for their behavioral dynamics score. For ELS animals, scores below zero indicated resilience while scores above zero indicated vulnerability. Although the control group showed comparable subdivisions, this aspect was not explored in the current analysis.

### 2.12 Interpretability analysis of behavior using supervised classification and SHAP

To investigate which features best discriminated between behavioral clusters, we trained a supervised classifier using pose estimation-derived features and evaluated its performance with interpretable machine learning techniques. We first aligned the DeepLabCut data with keypoint MoSeq-derived behavioral syllables and their corresponding behavioral clusters. Pose features were then extracted on a per-frame basis using egocentrically aligned coordinates, centering the animal’s body at the centroid and rotating it to align the tail-to-nose vector with the horizontal axis.

For each frame, we computed a feature set derived from the egocentrically aligned body posture. This included the two-dimension coordinates of all keypoints, pairwise Euclidean distances between body parts, the heading angle and angular velocity (based on the tail-to-nose vector) and the centroid position and velocity. To capture full-body orientation, we applied principal component analysis (PCA) to the set of body points and extracte the angle of the primary axis, along with its frame-to-frame angular velocity. Additional kinematic features included the change in heading angle over a fixed window (orientation change) and the displacement of the centroid over the same window. To encode temporal dynamics, we included lagged versions of all positional and angular features over a five-frame history and computed rolling statistics (mean, standard deviation, and sum) for all core spatial and kinematic features. This resulted in a complete spatiotemporal representation of posture and motion dynamics across time.

To classify behavioral clusters from these features, we trained an XGBoost model using a stratified 3-fold cross-validation scheme. The dataset was balanced across behavioral clusters by downsampling overrepresented classes to a maximum of 4000 randomly selected frames per cluster. Labels were numerically encoded using scikit-learn’s LabelEncoder and missing values were imputed using the column-wise mean. Model performance was evaluated via normalized confusion matrices, overall and per-class accuracy, and comparison to theoretical chance levels under uniform random assignment.

Model interpretability was assessed using SHAP (SHapley Additive exPlanations), a method based on cooperative game theory that quantifies the contribution of each feature to individual predictions^59^. We used the TreeExplainer to compute SHAP values for a randomly sampled 10% of the dataset on a final XGBoost model trained on the full data. SHAP summary plots were generated to visualize the most informative features globally (across all behaviors) and locally (within each cluster).

### 2.13 Statistical analysis

We used a mixed-effects linear model to assess the effect of condition on behavioral dynamic, frequency, transitions and bout duration scores. The model included random intercepts for animal and experiment to capture variability between animals and across experiments. Fixed effects included stress while random intercepts for experiment accounted for differences in behavioral responses across experiments. To test each behavioral cluster over time we used a mixed-effects linear model with fixed effects for stress, time and their interaction. Random intercepts for animal captured individual baseline differences in behavior and those for experiment accounted for potential variability across the two experiments. Model assumptions, including normality of residuals and homoscedasticity, were evaluated using Q–Q plots and residual versus fitted value plots. Although a model with experiment as a fixed effect showed a slightly lower Akaike information criterion (AIC), we preferred the original model because it better captured variability at both the animal and experiment levels. We consistently used this model except when the data showed collinearity.

For transition matrices and other discrete count data we opted for a GEE Negative Binomial model rather than a mixed-effects model. Model diagnostics included assessment of overdispersion through residual plots and evaluation of intra-cluster correlation. Given the overdispersion and clustering in our counts, the GEE framework allowed us to model the effect of condition on transition counts while clustering by experiment to account for within-experiment correlations.

## 3 Results

### 3.1 Automated freezing detection showed blunted freezing progression in ELS offspring

Our first goal was to establish a reliable, objective measure of freezing behavior using a supervised learning approach. Manual scoring is time consuming and prone to observer bias, so we used the SimBA framework^24^ to train a random forest classifier on hand-labeled freezing events. Automating the scoring facilitates consistent results across large datasets and reduces variability introduced by human scorers^60^.

We trained the classifier on a randomly selected subset of manually annotated video frames (labeled as freezing or not-freezing), along with features generated by SimBA from DeepLabCut pose estimation data. The full dataset included 182918 frames from 33 videos, using an 80/20 training/validation split. Once trained, the model was tested on an independent, held-out set of 211500 frames, where its predictions closely matched human scoring (r^2^ = 0.95, p < 0.001, **Fig. 3B**). This strong correlation supports using the classifier’s output as ground truth for all further analyses.

**Figure 3.**
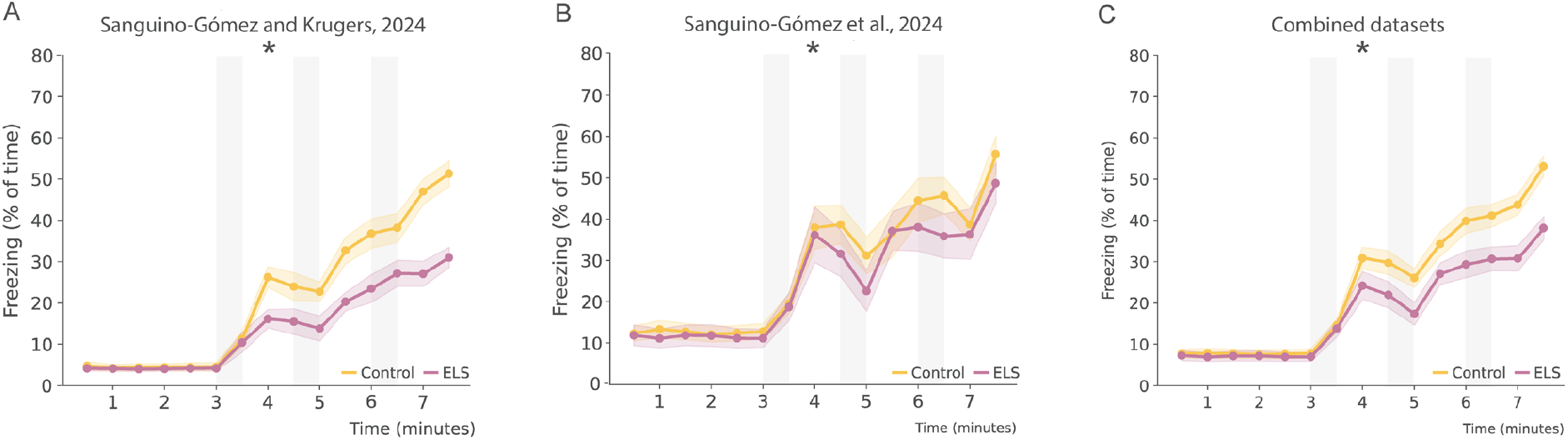
ELS reduces freezing behavior during FC across independent datasets. **A**. Lineplots showing freezing (mean % ± Standard error of the mean-SEM) across the course of conditioning in the Sanguino Gómez & Krugers (2024) dataset. ELS reduced freezing acquisition. **B**. Lineplots showing freezing (mean % ± SEM) during the FC task in the Sanguino Gómez et al. (2024) dataset. Freezing acquisition was decreased by ELS. **C**. Lineplots showing freezing (mean % ± SEM) over conditioning in the combined datasets. ELS reduced freezing acquisition. ELS blunted the freezing response. Shaded areas represent tone and footshock epochs. Sanguino Gómez & Krugers (2024) dataset: N_Control_ = 25, N_ELS_ = 25. Sanguino Gómez et al. (2024) dataset: N_Control_ = 16, N_ELS_ = 16. Combined datasets: N_Control_ = 41, N_ELS_ = 41. * ELS by time interaction effect. Effect p *≤* 0.05.

With our ground truth established, we next examined how a prior history of ELS affects the progression of freezing over time in the FC paradigm. To do this, we used two datasets generated in previous publications^46,47^, where this same classifier was used to score freezing behavior. For the current analysis, we focused only on control and ELS groups, excluding any additional manipulations to isolate the effects of ELS without interference from other variables (e.g. treatments).

Using a mixed linear model analysis, we confirm the main effect of time in the supervised data for both cohorts: Sanguino-Gómez and Krugers, 2024 (*β* = 3.721, SE = 0.128, z = 29.048, p < 0.001, **Fig. 3A**) and Sanguino-Gómez et al., 2024 (*β* = 4.412, SE = 0.191, z = 23.091, p < 0.001, **Fig. 3B**). A significant interaction between ELS and time was found (Sanguino-Gómez et al., 2024: *β* = −0.887, SE = 0.181, z = −4.898, p < 0.001, **Fig. 3A**; Sanguino-Gómez et al., 2024: *β* = −0.721, SE = 0.27, z = −2.667, p = 0.008, **Fig. 3B**), indicating that ELS animals showed a slower increase in freezing behavior over time when compared to controls (effect sizes: Cohen’s d = 0.284 and 0.189, respectively). A combined analysis confirmed the significant interaction between ELS and time (*β* = −0.822, SE = 0.154, z = −5.327, p < 0.001, Cohen’s d = 0.241, **Fig. 3C**).

Taken together, these findings show that our developed supervised classifier provided a reliable, scalable method for quantifying freezing behavior which highly correlates with human-anotated data from our datasets. Using this approach, we showed that ELS blunts the progression of fear-related freezing over time.

### 3.2 Unsupervised behavioral sequencing recapitulates the ELS effect on freezing found through supervised methods

After confirming that the supervised pipeline reliably scores freezing, we applied keypoint MoSeq^37^ to the same pose-estimation dataset to test whether the unsupervised motifs identified by keypoint MoSeq directly aligned to the manual freezing labels.

First, we trained keypoint MoSeq on the pose-estimation data, which generated 96 putative behavioral syllables (**Fig. 4A**). When matched to the supervised freezing labels, we found that a single dominant syllable (S0) accounted for roughly 75% of all freezing frames, leaving little room for contribution from other syllables to the overall recall (**Fig. 4A**). Given this imbalance, we set a precision threshold above 50% (**Fig. 4B**) to include an additional syllable (S28) that, despite low recall (**Fig. 4C**), contributed modestly to freezing behavior. Interestingly, we identified syllables with high precision for freezing that were exclusive to the ELS group and absent in controls (S61, S51 and S48 -**Fig. 4B**), suggesting subtle, ELS-specific differences in freezing behavior.

**Figure 4.**
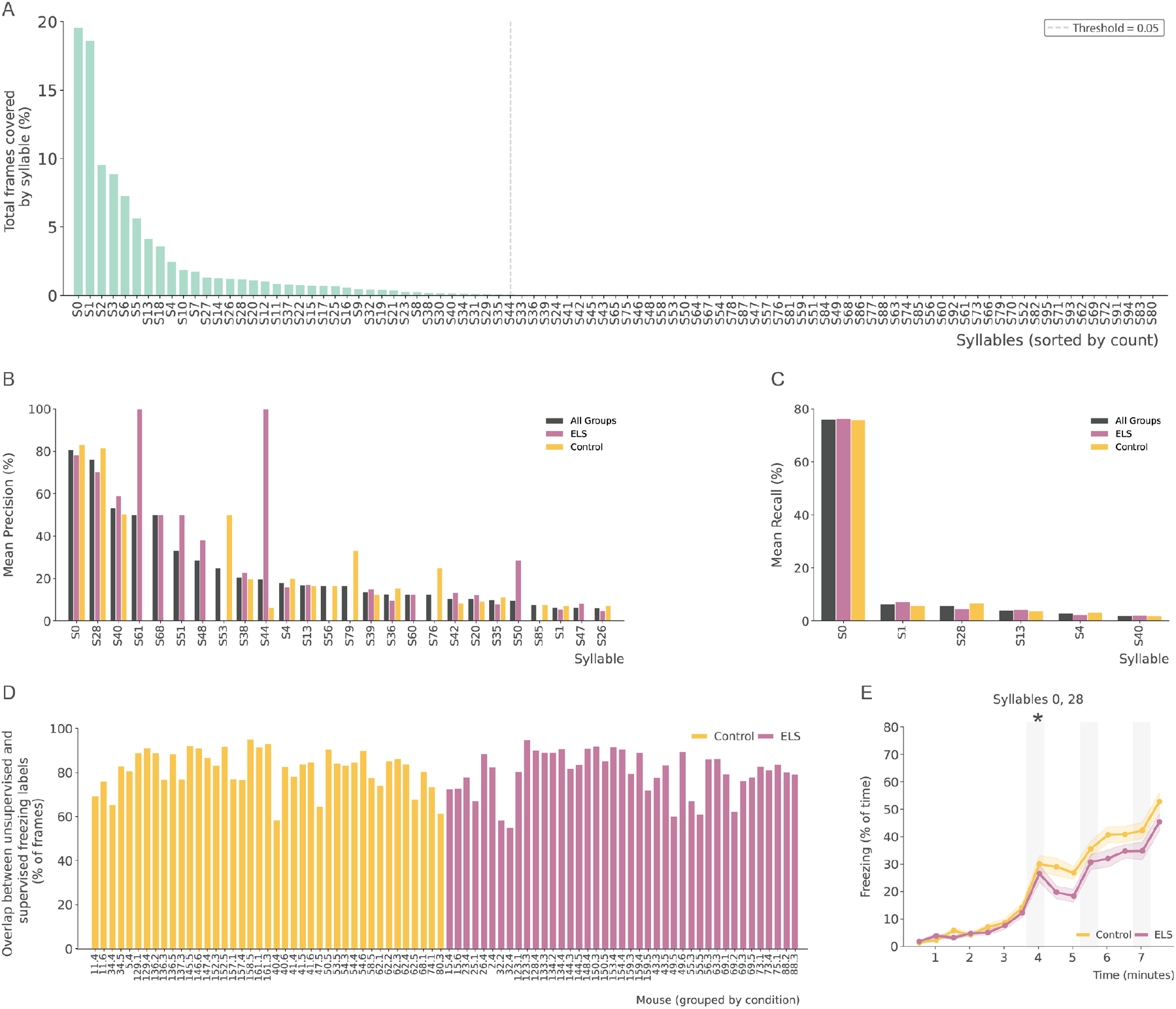
Validation of keypoint MoSeq-derived syllables using supervised freezing data from SimBA. **A**. Histogram of the percentage of frames covered by each keypoint MoSeq syllable across all animals. A 0.05% tail cutoff (dashed line) was applied to incorporate in our analysis the most frequently occurring syllables. **B**. Barplots of precision (mean % ± SEM) for each syllable versus supervised SimBA freezing annotations, shown combined but also separately for control and ELS groups. Syllables 0 and 28 show the highest precision, both within individual groups and when combined. **C**. Barplots of recall (mean % ± SEM) for each syllable against SimBA freezing labels in control and ELS animals combined and also individually analyzed. Syllable 0 contributed most to freezing annotations across groups and in the combined data. **D**. Barplots of overlap between SimBA freezing labels and syllables 0 and 28 (mean % ± SEM) for each animal. Every animal recapitulated over 50% of manual freezing annotations. **E**. Lineplots of syllable 0 and 28 usage (mean % ± SEM) over time during fear conditioning in control and ELS groups. ELS animals showed reduced use of these freezing associated syllables. Shaded areas represent tone and footshock epochs. N_Control_ = 41, N_ELS_ = 41. * ELS by time interaction effect. Effect p *≤* 0.05.

Some of these putative syllables, however, occurred with very low frequency. Therefore, we then ranked keypoint MoSeq-generated syllables by frequency and retained the smallest set that accounted for 99.5% of all syllable-labeled frames (**Fig. 2A**), leaving 38 syllables for downstream analysis.

To further validate the model, we combined the two surviving syllables (S0 and S28) that showed the highest freezing precision (**Fig. 4B**) and recall (**Fig. 4C**) and assessed their overlap with ground truth freezing labels from SimBA on a per-animal basis, stratified by group (**Fig. 4D**). Remarkably, the freezing detected by the supervised method overlapped with that identified by the unsupervised method overall on 80.86 ± 9.62% (Control: 80.12 ± 10.14%, ELS: 81.60 ± 9.01% -**Fig. 4D**). In all individual cases, the freezing coverage was well above 60% alignment with these syllables (**Fig. 4D**).

Next, we determined whether the ELS effect previously identified through supervised methods is replicated by using the unsupervised syllables defined by keypoint MoSeq. We analyzed the percentage of syllables S0 and S28 combined over time. The mixed linear model revealed a significant main effect of time (*β* = 0.126, SE = 0.004, z = 32.602, p < 0.001, **Fig. 4E**), indicating that the percentage of S0 + S28 syllables increased over time across all groups. Additionally, a significant interaction between ELS and time was observed (*β* = −0.023, SE = 0.005, z = −4.241, p < 0.001, **Fig. 4E**), suggesting that although both groups increased their use of S0 + S28 syllables, the rate of increase was slower in the ELS group compared to controls.

These findings consistently align with the ground truth data, demonstrating that unsupervised phenotyping effectively recapitulates the supervised results in freezing behavior.

### 3.3 Unsupervised behavioral sequencing methods reveals new behavioral motifs that differs in ELS animals

Once validated against known data, we used keypoint MoSeq to identify new sets of behaviors during FC and assess how a history of ELS alters their expression.

Syllables were visually inspected and grouped into ethologically-relevant clusters. Syllables that could not be correctly mapped because they encoded mixed behaviors or suffered from tracking inaccuracies were clustered separately and excluded from further analysis. On average, clusters corresponding to mixed behaviors covered 6.5% of the video frames, while those with inaccurate tracking accounted for 13.7%. Including unmapped frames, ≈ 20% of the video was not assigned to any syllable. The percentage of excluded data did not differ between stress conditions, except for inaccurate tracking, which was higher in control animals (*β* = −2.341, SE = 0.942, z = −2.486, p = 0.013 - **Supplementary Figure 1**).

Among the correctly mapped frames we identified seven ethologically relevant clusters: climbing, freezing, grooming, jump, locomotion, sniffing and turn (definitions in **Supplementary Table 1**). All behaviors occurred in all groups, in each experiment and in nearly all periods (**Supplementary material and methods**). When analyzing the overall frequency of behaviors, ELS animals exhibited fewer freezing events (*β* = −0.204, SE = 0.081, z = −2.511, p = 0.012, **Fig. 5A**) but more turn (*β* = 0.100, SE = 0.032, z = 3.099, p = 0.002, **Fig. 5A**) and sniffing (*β* = −0.068, SE = 0.185, z = −2.133, p = 0.033, **Fig. 5A**) events. When analyzed over time, some of these behaviors showed a significant ELS × time interaction effect. As shown by supervised methods^46,47^, unsupervised methods corroborate that the freezing cluster increased in both groups, with control animals showing a larger increase than ELS animals (*β* = −0.023, SE = 0.005, z = −4.241, p < 0.001, **Fig. 5B**). This method allowed us to further determine that ELS animals, relative to controls, began with a lower percentage of time spent in locomotion (*β* = 0.005, SE = 0.002, z = 2.553, p = 0.011, **Fig. 5C**) and a higher percentage in sniffing (*β* = −0.014, SE = 0.004, z = −3.656, p < 0.001, **Fig. 5E**) but these differences disappeared by the end of the trial. In contrast, turn behavior started similarly in both groups, but control animals showed a steeper decline over time (*β* = 0.026, SE = 0.007, z = 3.610, p < 0.001, **Fig. 5F**). There were no significant ELS × time interactions for grooming, locomotion or sniffing. The experimental data are reproducible across independent experiments, with the exception of locomotion measures **(Supplementary material and methods)**.

**Figure 5.**
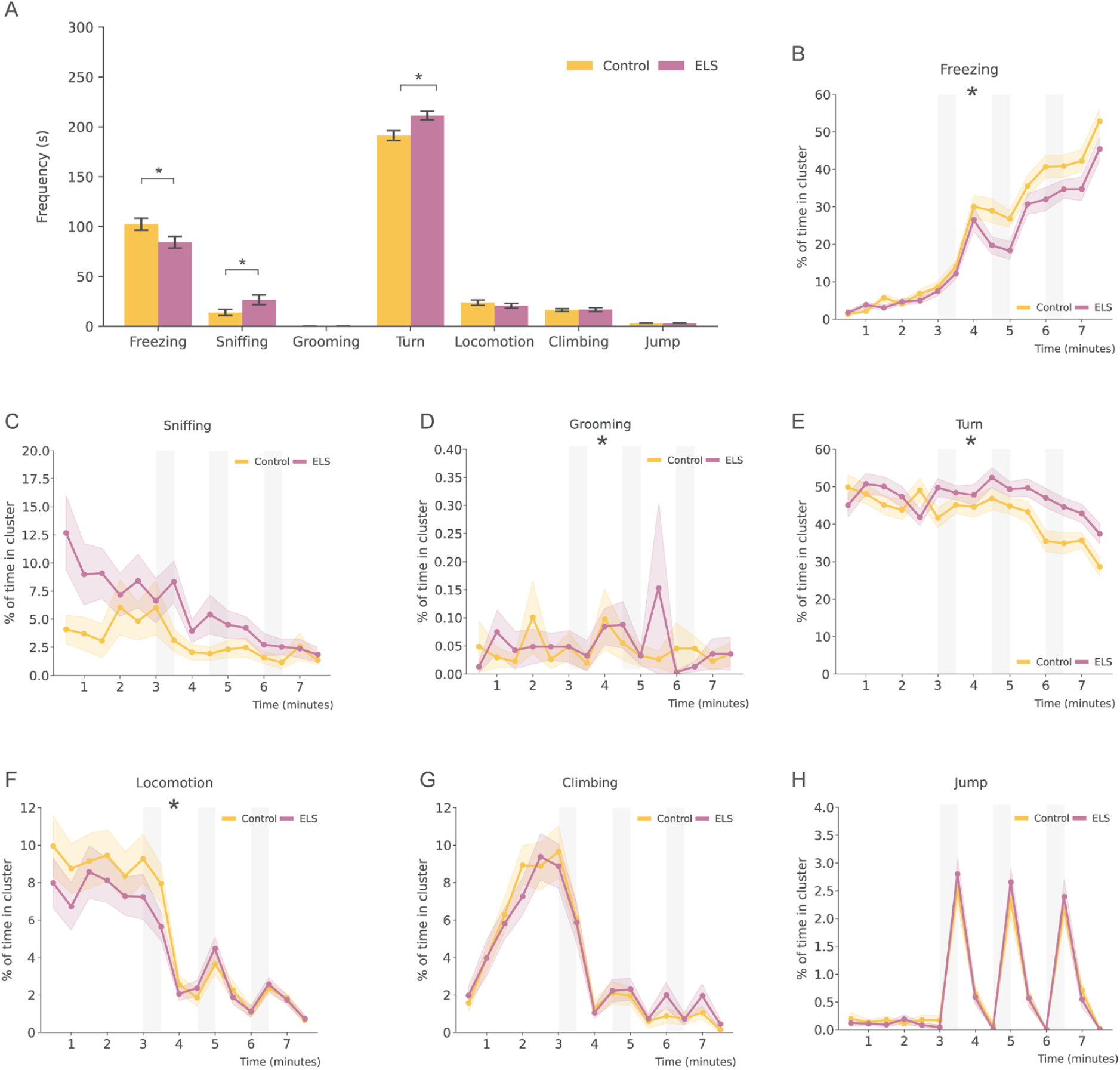
ELS alters keypoint MoSeq-derived behavior motifs during fear conditioning. **A**. Barplots showing total time in each behavioral cluster (mean in seconds ± SEM) across the entire FC session. ELS mice spent less time freezing and more time sniffing and turning compared to controls. **B**. Lineplots presenting freezing (mean % ± SEM) over time during FC in control and ELS animals. ELS mice showed a blunted freezing increase following shocks. **C**. Lineplots showing sniffing (mean % ± SEM) over time during FC in control and ELS animals. ELS animals began the session with elevated sniffing that normalized to control levels over time. **D**. Lineplots showing grooming (mean % ± SEM) over time during FC in control and ELS animals. No group differences were observed. **E**. Lineplots indicating turn (mean % ± SEM) over time during FC in control and ELS animals. Control animals showed a steeper reduction in turning after shocks compared to ELS mice. **F**. Lineplots showing locomotion (mean % ± SEM) over time during FC in control and ELS animals. ELS animals started with lower locomotion compared to controls. **G**. Lineplots showing climbing (mean % ± SEM) over time during FC in control and ELS animals. Climbing was unaffected by ELS. **H**. Lineplots showing jump (mean % ± SEM) over time during FC in control and ELS animals. Both groups showed similar jump spikes immediately after shocks. Shaded areas represent tone and footshock epochs. N_Control_ = 41, N_ELS_ = 41. * ELS effect (barplots) and ELS by time interaction effect (lineplots). Effect p *≤* 0.05.

Overall, these results suggest that ELS alters the course of several behaviors during the fear conditioning paradigm.

### 3.4 ELS reduces behavioral diversity in favor of prolonged active states

ELS may also alter the pattern in which behaviors occurred. To assess these changes, we plotted the incidence of behaviors over time and generated ethograms for each animal (see **Supplementary material and methods** for details and individual animals; the plot with the strongest correlation to the mean per condition is shown in **Fig. 6A** for Control and in **Fig. 6C** for ELS animals). We then identified the predominant behavior in each time bin to build a representative behavioral barcode, arranging behaviors from least to most active (see **Supplementary material and methods** for details and individual animals; the plot with the strongest correlation to the mean per condition is shown in **Fig. 6B** for Control and in **Fig. 6D** for ELS animals). This barcode serves as a quantitative snapshot of each mouse’s individual behavioral profile. Using this approach, we mapped all 82 animals and created representative videos for each condition (**Supplementary Material and Methods**).

**Figure 6.**
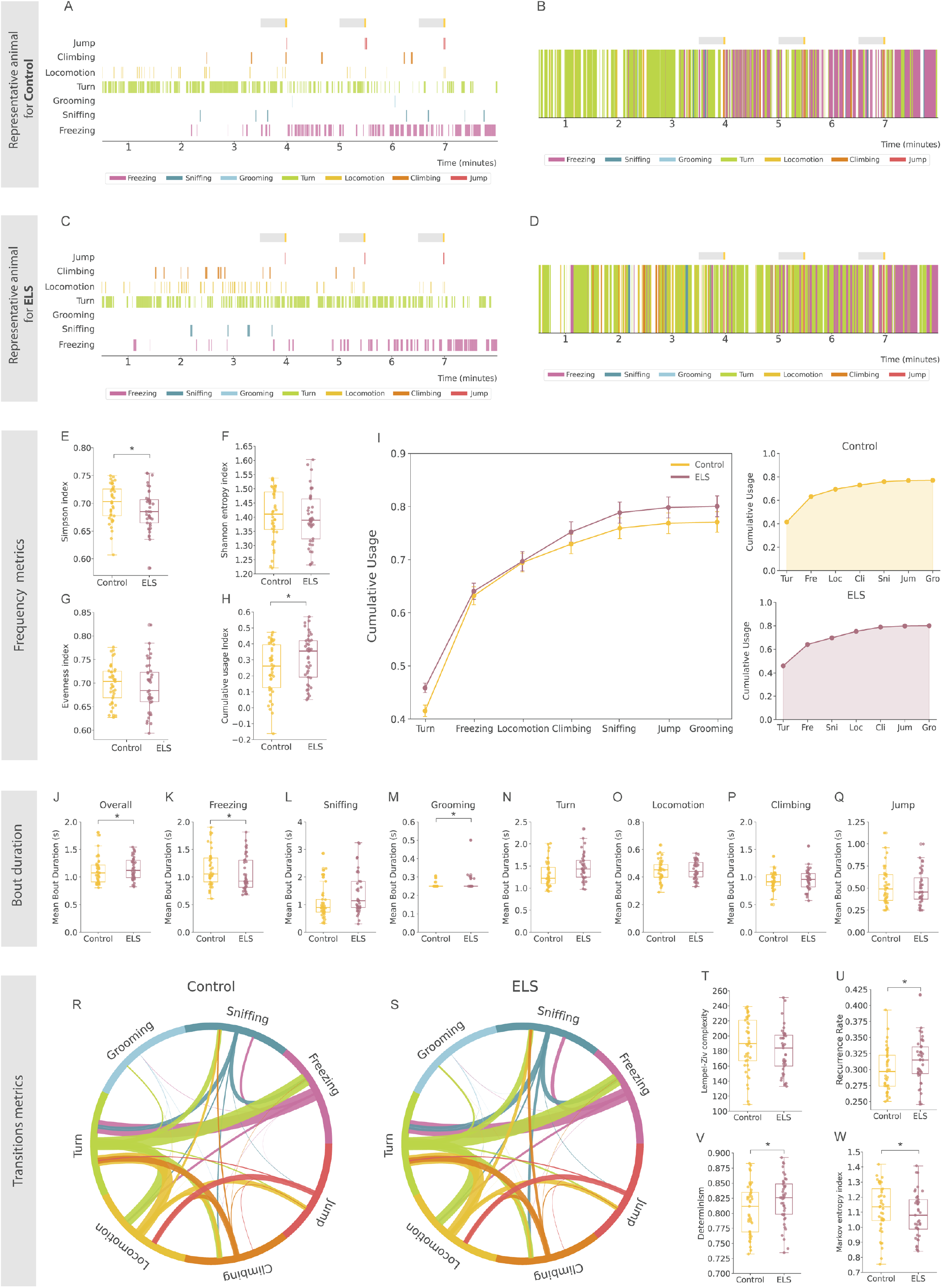
ELS reshapes diversity, structure, and transitions of keypoint MoSeq behavioral motifs during FC. **A**. Ethogram of the control mouse whose behavioral motif profile most closely matches the group average, shown with color-coded motifs at 250 ms resolution. Each row marks the active motif per frame. Gray and yellow bars indicate tone presentation and footshock delivery, respectively. **B**. Temporal barcode of the predominant motif per 250 ms bin for the same representative control mouse, highlighting motif prevalence across the full session. Gray and yellow bars indicate tone and footshock, respectively. **C**. Ethogram of the ELS mouse whose motif expression best approximates the group average, color-coded at 250 ms resolution. Gray and yellow bars mark tone and footshock presentation. ELS mice show fewer freezing ticks and more active motifs than controls. **D**. Temporal barcode of the predominant motif per 250 ms bin for the representative ELS mouse. Gray and yellow bars indicate tone and footshock, respectively. ELS animals show delayed freezing onset and sustained turning compared to controls. **E**. Boxplot of the Simpson diversity index (mean ± SEM). ELS reduces behavioral motif diversity. **F**. Boxplot of the Shannon diversity index (mean ± SEM). No difference between groups. **G**. Boxplot of the evenness index (mean ± SEM). Evenness is comparable across groups. **H**. Boxplot of the cumulative usage index (mean ± SEM). ELS increases overall motif usage. **I**. Linelot of cumulative motif usage (mean % ± SEM) ranked by total use. Left shows pooled curves, right shows each group separately. ELS alters the rank order of motif contributions. **J**. Boxplot of overall mean bout duration across all motifs (seconds ± SEM). No group differences. **K**. Boxplot of mean freezing bout duration (seconds ± SEM). ELS mice freeze in shorter bouts. **L**. Boxplot of mean sniffing bout duration (seconds ± SEM). Sniffing bouts are unchanged by ELS. **M**. Boxplot of mean grooming bout duration (seconds ± SEM). Grooming bouts are similar across groups. **N**. Boxplot of mean turn bout duration (seconds ± SEM). ELS mice turn in longer bouts. **O**. Boxplot of mean locomotion bout duration (seconds ± SEM). Locomotion bouts are unaffected by ELS. **P**. Boxplot of mean climbing bout duration (seconds ± SEM). Climbing bouts are unchanged by ELS. **Q**. Boxplot of mean jump bout duration (seconds ± SEM). Jump bouts are similar between groups. **R**. Chord diagram of motif-to-motif transition probabilities in controls. Arc width represents transition numbers and color shows starting behavioral motif of the transition. **S**. Chord diagram of transition probabilities in ELS mice. Arc width represents transition numbers and color shows starting behavioral motif of the transition. ELS alters overall the transition pattern in the FC task. **T**. Boxplot of Lempel-Ziv complexity index (mean ± SEM). No significant change between groups. **U**. Boxplot of recurrence rate (mean ± SEM). ELS increases recurrence. **V**. Boxplot of determinism index (mean ± SEM). ELS enhances determinism. **W**. Boxplot of Markov entropy index (mean ± SEM). ELS reduces Markov entropy. N_Control_ = 41, N_ELS_ = 41. * ELS effect. Effect p *≤* 0.05.

Using these ethograms, we showed that ELS animal showed a lower Simpson diversity score (*β* = −0.013, SE = 0.007, z = −2.039, p = 0.041, **Fig. 6E**), indicating that ELS offspring engaged in a narrower set of behaviors, with one or a few behaviors dominating their repertoire. Consistent with this observation, ELS animals also displayed a significantly higher CUI (*β* = −0.027, SE = 0.009, z = −3.079, p = 0.002, **Fig. 6H-I**), suggesting that the most frequently used clusters accounted for a larger share of their overall activity. No changes in frequency-based (Shannon) entropy or evenness (**Fig. 6F-G** respectively) were found, meaning that although the overall behavioral diversity decreased in ELS animals, the relative distribution and uniformity of behaviors remained similar across groups. Thus, ELS reduced behavioral diversity by limiting the repertoire but did not alter behavioral evenness or predictability.

Next, we computed the mean duration of the behavioral bouts. Overall, no differences were detected between groups; however, ELS animals exhibited distinct, cluster-specific changes. In particular, freezing bouts were notably shorter (*β* = −0.105, SE = 0.053, z = −2.001, p = 0.045, **Fig. 6J**), while more active behaviors, such as sniffing (*β* = 0.322, SE = 0.129, z = 2.506, p = 0.012, **Fig. 6K**) and turn(*β* = 0.159, SE = 0.052, z = 3.088, p = 0.002, **Fig. 6M**) have significantly prolonged bout durations.

Overall, ELS animals engage in a more restricted behavioral repertoire, characterized by prolonged and less variable active behaviors and a reduced duration of passive behaviors.

### 3.5 ELS reduces behavioral adaptability in favor of stereotypy

We next focused on the transitions between behaviors. The transition chorplots per group and the differences between groups are shown in **Fig. 6R** and **S**, respectively. We show no changes in Lempel ziv complexity (**Fig, 6T**) but an increase in the recurrence rate (*β* = 0.013, SE = 0.006, z = 2.264, p = 0.024, **Fig. 6U**) as well as in determinism (*β* = 0.016, SE = 0.005, z = 3.382, p = 0.001, **Fig. 6V**) meaning that even though the overall variety or richness in behavioral sequences stayed the same, ELS animals engage in more repetitive behaviors and behavioral sequences.

We also quantified entropy by analyzing the transition probabilities between distinct behavioral states using a first-order Markov chain model. ELS animals showed a significant reduction in Markov entropy (*β* = −0.041, SE = 0.018, z = −2.342, p = 0.019, **Fig. 6W**), suggesting that their behavioral sequences became more predictable and less variable. This reduced complexity may indicate a shift toward more stereotyped behaviors in the FC paradigm among the ELS offspring.

These results suggest that ELS animals show a more rigid and less flexible behavioral response in the FC task. Their behavior appears more repetitive and predictable, pointing to a shift toward stereotyped patterns that may reflect reduced adaptability when faced with a secondary stressor or challenge.

### 3.6 Individual multi-behavioral dynamics score allows to identify resilient profiles in ELS animals that behave similar to controls

For the subsequent analysis, we first binned the behavioral data by computing the relative frequency of each behavioral cluster in 30-second intervals and concatenated these values over time. Next, we built pairwise Euclidean distance matrices comparing every animal’s behavior. Then, we used MDS to map the resulting dissimilarity matrix into a two-dimensional map, preserving inter-animal distances by minimizing MDS’ stress function (^61,62^, **Fig. 2**)

Next, we computed a “behavioral dynamics score” defined as the log ratio of the Euclidean distance between an animal’s behavior and each group’s mean pattern (**Fig. 2**) in the MDS-defined space. This method parallels the behavioral flow score^36^ while incorporating a temporal component. Notably, ELS animals scored higher on this metric (*β* = 0.336, SE = 0.087, z = 3.869, p < 0.001, **Fig. 7B**).

**Figure 7.**
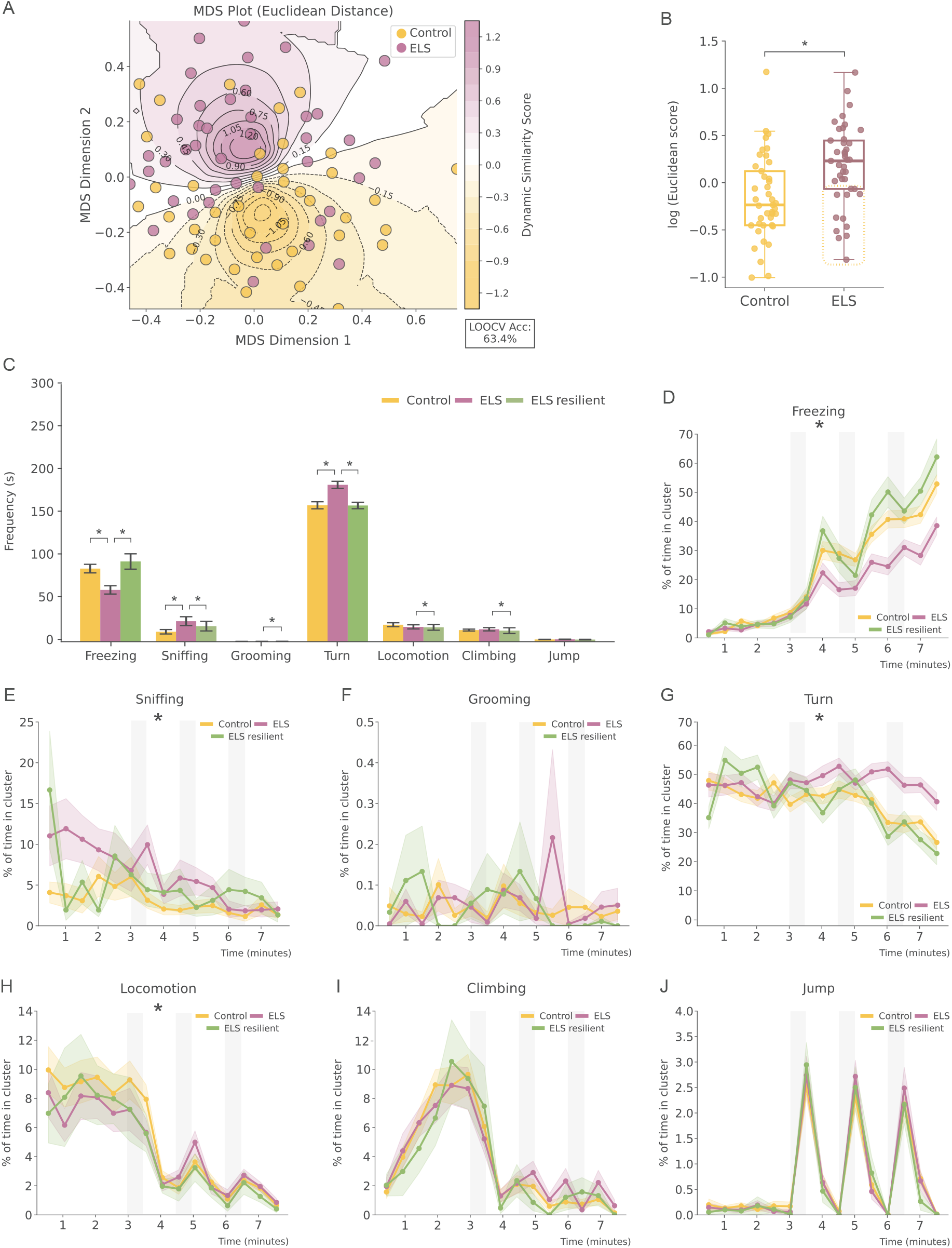
Behavioral dynamics allow identification of a resilient ELS subgroup with control-like motif frequency and dynamics. **A**. Multidimensional scaling (MDS) plot of the Euclidean distance matrix computed from pairwise comparisons of each animal’s behavioral-dynamics profiles. Each dot is the two-dimensional embedding of one mouse, positioned so that the distance between dots reflects the original Euclidean distances in feature space. Shaded ellipses represent the group median (dark core = median, fading edges = dispersion). Leave-one-out cross-validation accuracy (LOOCV Acc) is indicated in the bottom corner. **B**. Boxplots of the Euclidean scores (mean ± SEM) for control versus ELS animals. ELS mice differed significantly from the control animals. Enclosed with a square are represented the ELS animals with scores below zero and therefore classified as resilient. **C**. Barplots showing total time in each behavioral cluster (mean in seconds ± SEM) across the entire FC session across control, ELS and ELS resilient groups. ELS resilient mice spend more time freezing and less time sniffing and turning than ELS animals, matching control levels. **D**. Lineplots presenting freezing (mean % ± SEM) over time during FC in control, ELS and ELS resilient animals. Resilient mice override the ELS-driven freezing deficit and match or surpass control freezing levels. **E**. Lineplots presenting sniffing (mean % ± SEM) over time during FC in control, ELS and ELS resilient animals. ELS mice show elevated early sniffing that normalizes to control levels in resilient animals. **F**. Lineplots showing grooming (mean % ± SEM) over time during FC in control, ELS and ELS resilient animals. Grooming dynamics are comparable across groups. **G**. Lineplots showing turn (mean % ± SEM) over time during FC in control, ELS and ELS resilient animals. ELS resilient mice suppress turning during footshocks similarly to controls. **H**. Lineplots showcasing locomotion (mean % ± SEM) over time during FC in control, ELS and ELS resilient animals. ELS mice show locomotion suppression during shocks that is absent in resilient animals. **I**. Lineplots presenting climbing (mean % ± SEM) over time during FC in control, ELS and ELS resilient animals. Climbing remains stable across conditions. **J**. Lineplots showing jump (mean % ± SEM) over time during FC in control, ELS and ELS resilient animals. All groups display similar jump peaks following shocks. Shaded areas represent tone and footshock epochs. N_Control_ = 41, N_ELS_ = 29, N_ELS resilient_ = 12. * ELS effect (barplots) and ELS by time interaction effect (lineplots). Effect p *≤* 0.05.

To verify this approach, we also computed (dis)similarity matrices using several other metrics-including correlation, cosine, Jensen–Shannon, and Manhattan distances- to assess result consistency (see **Supplementary Figure 3** and (**Supplementary Materials and Methods**). After testing several metrics, we selected the Euclidean distance score for further analysis because it achieved relatively high leave-one-out cross-validation (63.5%, **Fig. 7A**).

In parallel, we also computed similarity scores based on transition probabilities, following the method in von Ziegler et al. 2024, rather than using temporal dynamics. This transition-based approach showed an overlap of 58.3% with the dynamics-based resilient classifications **Supplementary Materials and Methods**), providing further support for the robustness of the dynamics score used here.

Within the ELS group, animals with behavioral dynamics scores below zero-indicating behaviors similar to controls-were classified as “resilient.” Out of 41 ELS animals, 12 met this criterion (6 from^46^ and 6 from^47^), which accounts for ≈ 30% of the ELS population. The overlap of resilient classifications between the Euclidean method and other metrics ranged from 60% and 100%, supporting its use as our resilience criterion (**Supplementary Materials and Methods**).

Using this new classification (control, ELS and resilient animals) we then reexamined the frequency of each behavior across groups. Resilient animals froze more than vulnerable ELS (*β* = 0.180, SE = 0.038, z = 4.681, p < 0.001, **Fig. 7C**) and even surpassed the freezing levels observed in controls (*β* = −0.342, SE = 0.082, z = −4.168, p < 0.001, **Fig. 7C**). These animals also showed reduced sniffing (*β* = 0.492, SE = 0.119, z = 4.139, p < 0.001, **Fig. 7C**), grooming *β* = −0.398, SE = 0.187, z = −2.123, p = 0.003, **Fig. 7C**), turn (*β* = 0.009, SE = 0.016, z = 0.568, p < 0.001, **Fig. 7C**), locomotion (*β* = −0.424, SE = 0.089, z = −4.755, p < 0.001, **Fig. 7C**) and climbing (*β* = −0.196, SE = 0.074, z = −2.624, p = 0.009, **Fig. 7C**) compared with the typical ELS offspring.

Over time, resilient animals exhibited a steeper increase in freezing behavior relative to ELS (*β* = −1.865, SE = 0.249, z = −7.484, p < 0.001, **Fig. 7D**) and even steeper than controls animals (*β* = 0.619, SE = 0.238, z = 2.599, p = 0.009, **Fig. 7D**). They also showed decreased turn over time compared with ELS (*β* = 1.622, SE = 0.326, z = 4.967, p < 0.001, **Fig. 7E**) and they started with marginally less sniffing relative to ELS (*β* = −0.338, SE = 0.176, z = −1.920, p = 0.05, **Fig. 7G**).

Representative ethograms and barplots showing the animal with the strongest correlation with the mean for each newly defined condition are presented in **Fig. 8A, C** and **E**, for ethograms, and in **Fig. 8B, D** and **F**, for boxplots, respectively. Overall, differences between control and ELS animals were mainly driven by the vulnerable subgroup. For instance, vulnerable ELS animals showed lower Simpson diversity than controls (*β* = − 0.022, SE = 0.008, z = − 2.835, p = 0.005, **Fig. 8G**). And within the ELS cohort, resilient animals resembled controls, while vulnerable animals exhibited a more limited behavioral repertoire than the resilient subgroup (*β* = 0.029, SE = 0.011, z = 2.627, p = 0.009, **Fig. 8G**). Likewise, the higher CUI observed in ELS animals relative to controls comes solely from the vulnerable subgroup (*β* = 0.075, SE = 0.028, z = 2.664, p = 0.008, **Fig. 8J**). No changes in Shannon entropy or evenness were found (**Fig. 8H** and **I**).

**Figure 8.**
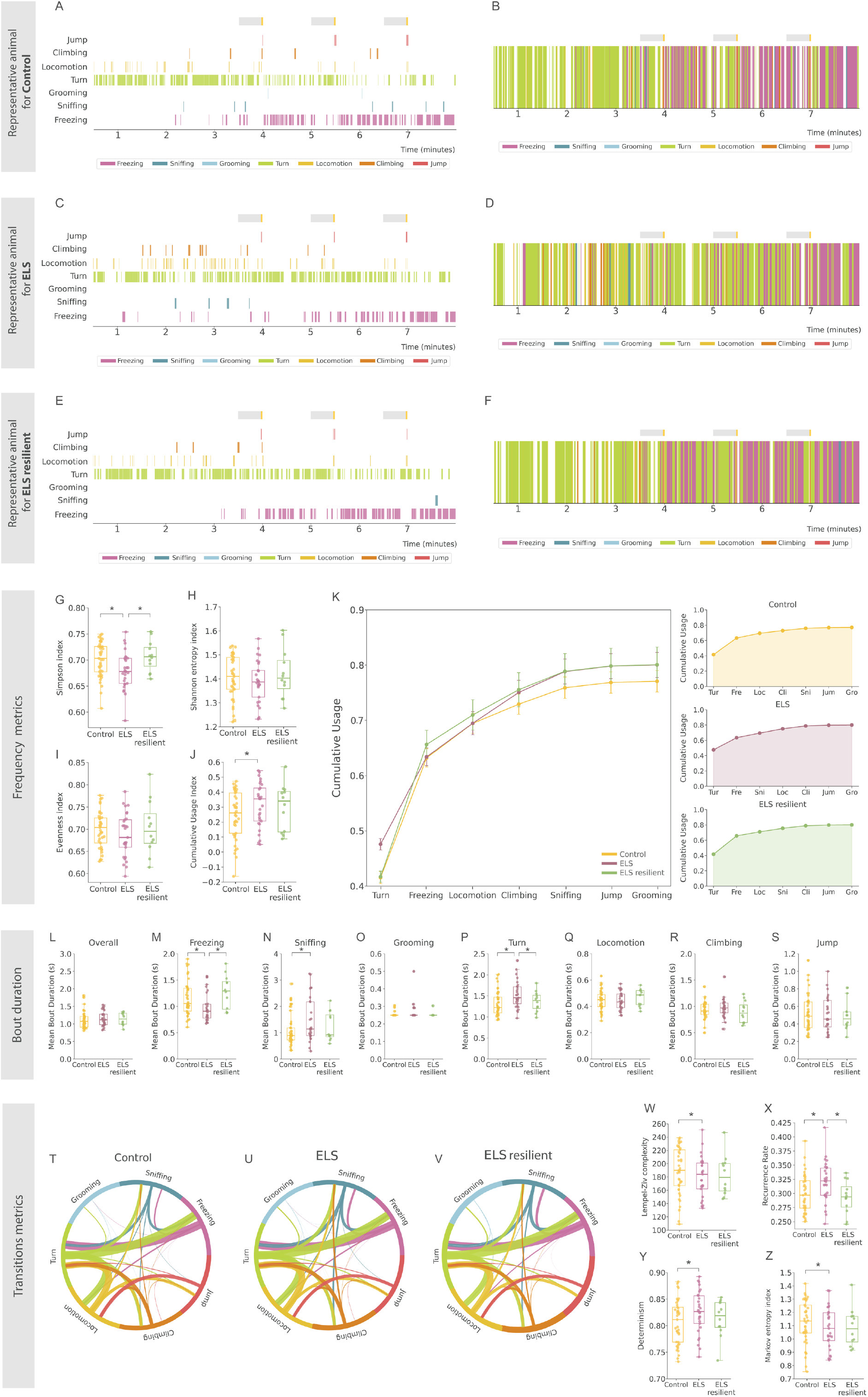
ELS resilient mice recapitulate control-like motif characteristics and transition dynamics during FC. **A**. Ethogram of the control mouse whose behavioral motif profile most closely matches the group average, shown with color-coded motifs at 250 ms resolution. Each row marks the active motif per frame. Gray and yellow bars indicate tone presentation and footshock delivery, respectively. **B**. Temporal barcode of the predominant motif per 250 ms bin for the same representative control mouse, highlighting motif prevalence across the full session. Gray and yellow bars indicate tone and footshock, respectively. **C**. Ethogram of the ELS mouse whose motif expression best approximates the group average, color-coded at 250 ms resolution. Gray and yellow bars mark tone and footshock presentation. ELS mice show fewer freezing ticks and more active motifs than controls. **D**. Temporal barcode of the predominant motif per 250 ms bin for the representative ELS mouse. Gray and yellow bars indicate tone and footshock, respectively. **E**. Ethogram of the ELS resilient mouse whose motif expression best approximates the group average, color-coded at 250 ms resolution. Gray and yellow bars mark tone and footshock presentation. ELS resilient mice mirror control motif patterns. **F**. Temporal barcode of the predominant motif per 250 ms bin for the representative ELS resilient mouse. Gray and yellow bars indicate tone and footshock, respectively. ELS resilient mice show more freezing ticks and less active motifs than ELS, matching control-like patterns. **G**. Boxplot of the Simpson diversity index (mean ± SEM) by group. ELS reduces behavioral motif diversity, while resilient mice show a recovery to control-like levels. **H**. Boxplot of the Shannon entropy index (mean ± SEM). No significant group differences. **I**. Boxplot of the evenness index (mean ± SEM). Evenness is comparable across all groups. **J**. Boxplot of the cumulative usage index (mean ± SEM). ELS resilient mice restore overall motif usage to levels observed in controls. **K**. Lineplots of cumulative motif usage (mean % ± SEM) ranked by total use. Left panel shows pooled curves, right panels show each group separately. Resilient mice shift rank order almost entirely back toward control levels. **L**. Boxplot of overall mean bout duration across all motifs (seconds ± SEM). There are no group differences. **M**. Boxplot of mean freezing bout duration (seconds ± SEM). Resilient mice freeze in longer bouts than ELS, matching controls. **N**. Boxplot of mean sniffing bout duration (seconds ± SEM). Sniffing bouts are lower in ELS animals but they don’t significantly return to control-like duration. **O**. Boxplot of mean grooming bout duration (seconds ± SEM). Grooming bouts are similar across groups. **P**. Boxplot of mean turn bout duration (seconds ± SEM). Resilient mice turn in shorter bouts than ELS, similar to controls. **Q**. Boxplot of mean locomotion bout duration (seconds ± SEM). Locomotion bouts are unaffected by ELS or resilience. **R**. Boxplot of mean climbing bout duration (seconds ± SEM). Climbing bouts are unchanged across groups. **S**. Boxplot of mean jump bout duration (s ± SEM). Jump bouts are similar across all groups. **T**. Chord diagram of motif-to-motif transition probabilities in controls. Arc width represents transition numbers and color shows starting behavioral motif of the transition. **U**. Chord diagram of motif-to-motif transition probabilities in ELS animals. Arc width represents transition numbers and color shows starting behavioral motif of the transition. ELS animals have different transition dynamics in the FC task. **V**. Chord diagram of motif-to-motif transition probabilities in ELS resilient animals. Arc width represents transition numbers and color shows starting behavioral motif of the transition. Transition dynamics in the FC task differ from those of control and vulnerable ELS groups. **W**. Boxplot of the Lempel-Ziv complexity index (mean ± SEM). When subsetted, ELS-vulnerable mice show reduced behavioral complexity compared to controls. **X**. Boxplot of recurrence rate (mean ± SEM). ELS increases recurrence, but resilient mice display recurrence levels comparable to controls. **Y**. Boxplot of determinism index (mean ± SEM). ELS enhances determinism but ELS resilient mice normalize to control levels. **Z**. Boxplot of Markov entropy index (mean ± SEM). Once separated, ELS vulnerable animals have lower Markov entropy compared to controls. N_Control_ = 41, N_ELS_ = 29, N_ELS resilient_ = 12. * ELS effect. Effect p *≤* 0.05.

Within the ELS subgroups, resilient animals not only froze more but also for longer durations compared to vulnerable ELS (*β* = 0.263, SE = 0.089, z = 2.969, p = 0.003, **Fig. 6L**). They also turned less frequently and, when turning, their turn bouts were shorter (*β* = −0.201, SE = 0.089, z = −2.255, p = 0.024, **Fig. 8O**). Although sniffing behavior did not differ significantly between the ELS subgroups, vulnerable animals spent sniffing longer periods than controls (*β* = −0.420, SE = 0.128, z = −3.285, p = 0.001, **Fig. 8M**), while resilient animals did not differ significantly from either group.

We then recalculated the behavioral transitions for the three designated categories (differential chordplots between conditions **Fig. 8T-V**). Once substratified, vulnerable ELS animals maintained a significant increase in recurrence rate (*β* = 0.022, SE = 0.008, z = 2.798, p = 0.005, **Fig. 8X**) and determinism (*β* = 0.022, SE = 0.008, z = 2.803, p = 0.005, **Fig. 6Y**) and they also showed now a reduction in Lempel–Ziv complexity (*β* = −10.590, SE = 5.186, z = −2.042, p = 0.041, **Fig. 8W**) compared with controls. Resilient animals generally had intermediate scores; however, in recurrence rate they were closer to controls and significantly different from the vulnerable subgroup (*β* = −0.029, SE = 0.011, z = −2.727, p = 0.006, **Fig. 8X**). Finally, vulnerable ELS animals exhibited increased Markov entropy (*β* = −0.051, SE = 0.026, z = −1.951, p = 0.050, **Fig. 8Z**) compared to controls.

In summary, by integrating multi-behavioral temporal dynamic patterns discovered through unsupervised methods, we identified a resilient subpopulation within the ELS group with distinct behavioral trajectories compared to typical ELS subjects. Resilient animals showed prolonged passive behavior and shorter active responses, resulting in a more diverse, flexible and unpredictable behavior, diverging from the stereotypical pattern characteristic of the ELS offspring and aligning more with the control condition.

### 3.7 Understanding behavioral clusters using interpretable machine learning

To assess how well keypoint MoSeq-identified behaviors could be distinguished based on pose dynamics, we trained a supervised classifier to predict the behavioral clusters from pose-derived features. The DeepLabCut data was egocentrically aligned and from it, we extracted a comprehensive set of features, including two-dimensional keypoint positions, pairwise distances between body parts, heading angles, angular velocity, global body orientation (estimated via PCA) and both individual body parts and centroidal velocities (**Supplementary material and methods**). Temporal structure was retained using rolling window statistics (mean, standard deviation, and sum) and time-lagged features across a 5-frame window.

With those features, we trained a gradient-boosted decision tree model (XGBoost) with 3-fold stratified cross-validation and evaluated its performance relative to chance. The classifier reached an overall accuracy of 62.7%, substantially above the chance level of 12.5%-an absolute improvement of 50.2% and a 5.0*×* gain over random guessing (**Fig. 9A**).

**Figure 9.**
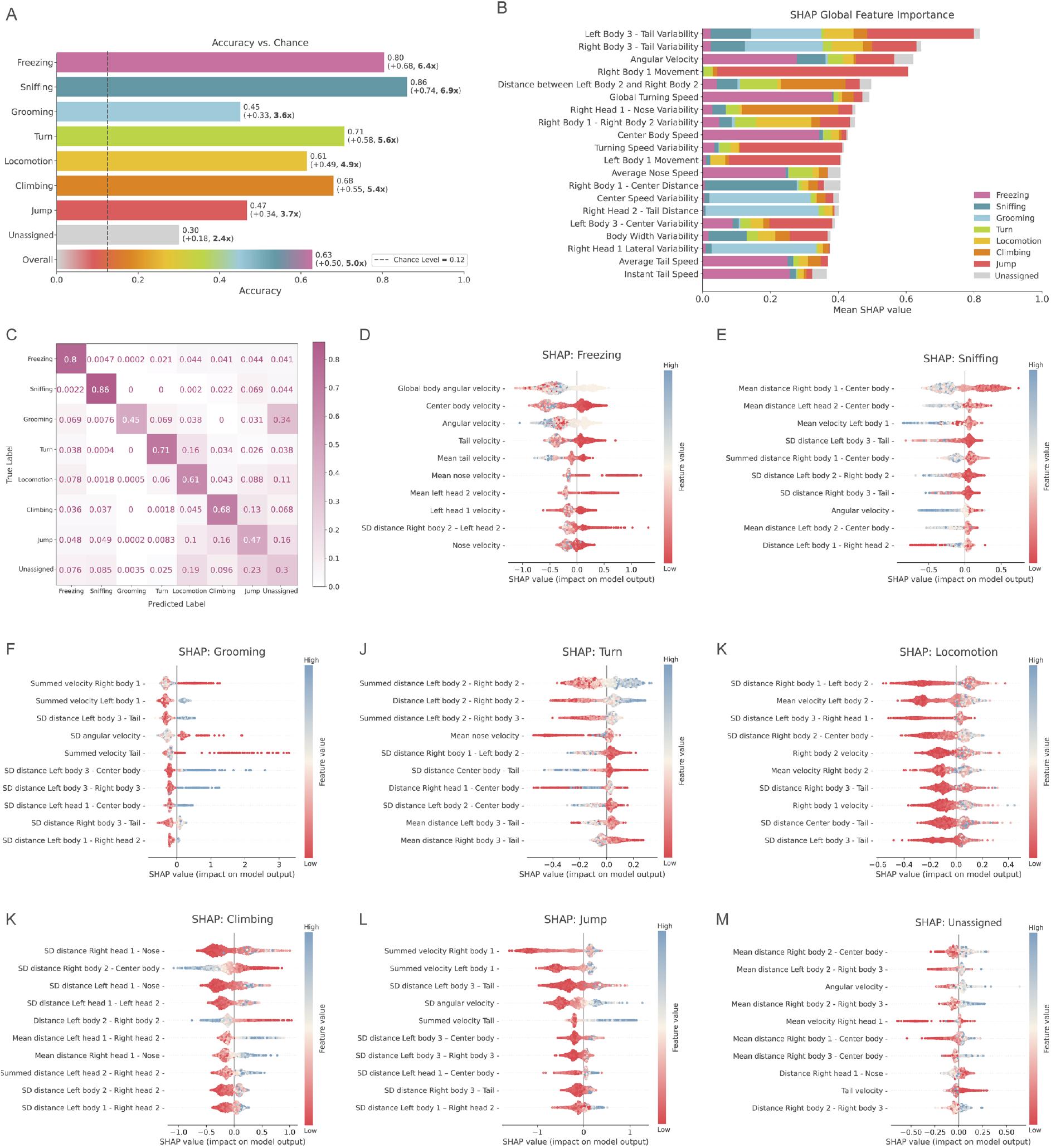
SHAP based explainability shows kinematic feature contributions to the unsupervised behavior classification. **A**. Barplots of classification accuracy (mean ± SEM) for each behavior, with the dashed line marking chance level (0.12). **B**. Global SHAP feature-importance bar chart showing SHAP values (mean ± SEM) for the top 20 kinematic metrics, with bar segments color-coded by the behavior they most strongly predict. **C**. Normalized confusion matrix showing the fraction of true versus predicted labels across the eight behavior classes. **D**. SHAP violin plot for freezing predictions, showing the distribution of SHAP values (mean ± SEM) for the top 10 features impacting freezing classification. **E**. SHAP violin plot for sniffing predictions, showing the distribution of SHAP values (mean ± SEM) for the top 10 features impacting sniffing classification. **F**. SHAP violin plot for grooming predictions, showing the distribution of SHAP values (mean ± SEM) for the top 10 features impacting grooming classification. **G**. SHAP violin plot for turn predictions, showing the distribution of SHAP values (mean ± SEM) for the top 10 features impacting turn classification. **H**. SHAP violin plot for locomotion predictions, showing the distribution of SHAP values (mean ± SEM) for the top 10 features impacting locomotion classification. **I**. SHAP violin plot for climbing predictions, showing the distribution of SHAP values (mean ± SEM) for the top 10 features impacting climbing classification. **J**. SHAP violin plot for jump predictions, showing the distribution of SHAP values (mean ± SEM) for the top 10 features impacting jump classification. **K**. SHAP violin plot for unassigned predictions, showing the distribution of SHAP values (mean ± SEM) for the top 10 features impacting theunassigned class.

Classification accuracy varied across behavioral motifs. Freezing and climbing were predicted with high accuracy (86.1% and 80.4%, respectively), followed by jump (70.6%), sniffing (67.9%) and locomotion (61.5%) (**Fig. 9A**). In contrast, behaviors like grooming (45.0%), turn (46.7%) and “unassigned” (30.1%) (**Fig. 9A**) were more difficult to classify accurately. A normalized confusion matrix (**Fig. 9C**) shows that misclassifications predominantly occur among the more movement-rich motifs (e.g. grooming is often mistaken for unlabeled behavior, and turning is occasionally defined as locomotion). These results suggest that while some motifs are clearly distinguisable based on pose dynamics alone, others may require additional contextual or temporal information for accurate disambiguation.

To understand which features contributed most to the classification, we used SHAP (SHapley Additive exPlanations) to interpret the model’s predictions. Global SHAP scores identified that variability in body tilt, angular velocity and global turning speed contributed the most to class discrimination across clusters (**Fig. 9B**). Importantly, feature importance varied by cluster, reflecting distinct pose dynamics associated with different behaviors (**Fig. 9D-K**).

Freezing (**Fig. 9D**) is characterized by near-zero centroid speed and minimal lean variance with brief angular-velocity spikes. Sniffing (**Fig. 9E**) relies on distinct head-to-body centroid distances. Grooming (**Fig. 9F**) shows moderate summed limb and head velocities that reflect localized repetitive motion. Turning (**Fig. 9G**) involves increased separation of left-side body points related to the hunched posture. Locomotion (**Fig. 9H**) depends on rhythmic swings in limb-to-center distances and steady mean velocities. Climbing (**Fig. 9I**) displays high variance in head-to-body distance and body lean. Jump (**Fig. 9J**) is defined by sharp peaks in summed velocity and angular acceleration. Unassigned frames (**Fig. 9K**) produce weak, dispersed feature contributions consistent with their low accuracy. These explainability features align with the behavioral predefinitions (**Supplementary Table 1**) that were used to classify keypoint MoSeq–generated syllables into behavioral clusters through visual inspection, therefore validating such classification.

In conclusion, we demonstrate that pose estimation-derived features can confidently predict keypoint MoSeq behavioral clusters with high accuracy and that interpretable machine learning tools like SHAP can clarify the underlying structure of the unsupervised-generated behavioral motifs.

## 4 Discussion

This study integrates supervised and unsupervised methodologies to examine how ELS shapes multi-dimensional behavioural dynamics during a FC task. First, we used a supervised classifier to reliably detect freezing behavior and found that adult male mice that were exposed to early life adversity between PND 2-9 showed a slower increase in freezing behavior over the course of three tone-footshock associations, when compared to controls. We then validated these results with unsupervised clustering, which not only confirmed the differences in freezing behavior but also revealed additional behaviors that were altered by ELS. By analyzing the frequency and transition dynamics between behavioral states, we observed that ELS animals tend to engage in more narrow, rigid, repetitive, predictable and stereotyped behavioral patterns, characterized for more frequent and longer active behaviors. This detailed behavioral mapping further allowed us to identify a resilient subgroup among the ELS animals, characterized by distinct behavioral trajectories that were comparable to or sometimes even surpassed behavior in control mice, lower entropy, greater behavioral diversity and enhanced frequency and duration of passive behaviors.

Unsupervised clustering through keypoint MoSeq showed that what we broadly manually labeled as freezing behavior may actually consist of two distinct subtypes of behaviors. Visual inspection indicated that cluster freezing comprises two behavioral syllables: S0, which resembles a classic, elongated freezing posture, and S28, which is characterized by a more curled and compactposture resembling a huddling-like form. Huddling has been described as a protective behavior known as defensive aggregation, where social animals such as rodents cluster together in response to threats like predators^63^. While huddling is typically a group behavior, its appearance in isolated animals may reflect an instinctive attempt to self-soothe or reduce perceived exposure to threat^63,64,65^. In our case, S28 predominantly occurred near the walls of the arena, suggesting a compensatory behavior, seeking tactile contact with the environment to mimic the comfort of conspecific proximity^63^. The observed behavioral distinction implies that freezing is not a monolithic response but rather a set of functionally distinct passive coping strategies. While both S0 and S28 reflect immobility, the latter’s curled posture and reduced body extension may signal a defensive effort to protect vital organs, a state of overwhelming stress^63,66,67^ or even learned helplessness -a response observed when animals in uncontrollable threat contexts, stop attempting to escape and prepares for an inevitable outcome^68^.

Additionally, the supervised classifier detected three other “freezing” syllables (S51, S61 and S68) exclusively present in the ELS animals, with a precision over the supervised labels above 50%. Due to their low occurrence, these behaviors were excluded from the main analysis but their unique presence in ELS animales may indicate subtle differences in freezing response after a previous history of stress that standard freezing analysis may have overlooked.

Beyond freezing, our clustering analysis revealed several etiologically relevant behavioral patterns that, from more passive to more active were labelled as: freezing, sniffing, grooming, turn, locomotion, climbing and jump.

First, it is important to note that 20% of the frames didn’t correspond to any specific behavior. Those frames were therefore left uncategorized. This came from errors in the pose estimation data or that, due to our strict definitions, syllables couldn’t fit in any of the categories presented above; because there was no clear indication of a specific behavior displayed or because the syllable represented a mix of behaviors. We also acknowledge that using a single top-view camera limited our ability to see certain body parts (e.g. limbs), which may have provided more insight from our pose estimation data and helped broaden our behavioral repertoire. We faced challenges to define behaviors that from a top view could be interpreted by the keypoint MoSeq as similar (e.g. turn and climbing) and therefore post-hoc scripts were developed to for instance, assess the distance to the walls to be able to discriminate between both scenarios.

Although most behaviors were observed in both conditions, ELS animals showed lower frequency and shorter duration of freezing events overall and an increase in frequency and duration of more active behaviors like turning and sniffing. Over the course of training in the FC paradigm, control animals tended to shift more quickly toward a predominantly freezing (passive) strategy, whereas ELS animals exhibited a slower transition marked by sustained levels of exploratory and dynamic behaviors during the stressful task. These data corroborate our own results obtained in a supervised manner^46,47^ and others^49,50^. Interestingly, Bordes et al. (2024) also used unsupervised behavioral methods to further analyze behavioral response of ELS mice during FC. For that, they used a combination of DeepLabCut and their own package DeepOF^69,27^ and they found no changes in behavior in ELS males -in concordance to^70^- but a decrease in passive behaviors and more active, explorative behaviors in females^49^, comparable to our findings in male mice

These results may reflect a broader alteration in coping strategies in response to a stressful learning paradigm, where a history of ELS leads to prolonged engagement in active behaviors at the expense of passive responses. Given that active and passive coping might rely on distinct brain circuits (active coping: amygdala–nucleus accumbens; passive coping: amygdala–periaqueductal gray;^71,72^) it is possible those networks are selectively affected by early stressful experiences.

Although freezing is traditionally considered the primary fear response in the FC paradigm, growing evidence suggests that conditioned cues also trigger a range of active behaviors: running, rearing, jumping, or suppressing ongoing reward-seeking^43,73,74,75^. These responses often follow a temporal pattern: active behaviors emerge early during the cue presentation, while freezing tends to appear later in time as shock becomes more likely^71^. Still, the field has historically focused on freezing; likely because of experimental setups that bias toward its expression (e.g. small spaces, auditory cues, intense shocks), its ease of measurement and its long-standing conceptual dominance^45^, thereby limiting recognition of the full behavioral repertoire elicited during the FC task.

Our findings add to this broader perspective. In our studies, ELS animals did not just freeze less; they showed a wider, more structured behavioral repertoire during cue exposure. These were not random or chaotic responses, but represented repeated patterns that may point to different coping within a fear context^10^. This suggests that the range and structure of behaviors-not just freezing-is relevant when studying fear conditioning and how this repertoire is modulated by, for example, a history of ELS.

In addition, freezing behavior itself is increasingly seen not as a passive shutdown but as a complex cognitive state that enables threat assessment and action preparation^76,77^. Far from behavioral shutdown, freezing can reflect an orchestrated inhibition of movement involving parasympathetic dominance and coordinated brain activity across the amygdala, prefrontal cortex, and periaqueductal gray^76,77^, optimizing internal conditions for decision-making. Thus, a reduction or delay in freezing responses, such as the currently observed in ELS animals, may indicate not just altered motor output but also an impaired capacity for adaptive threat assessment. In clinical contexts, altered freezing behavior has been associated with PTSD and anxiety disorders^78^, reinforcing its importance as a flexible, preparatory process rather than only a fear index. Interestingly, our barcodes suggest that freezing in ELS animals emerges later in the FC task, indicating a delayed response rather than a complete absence of this behavior. This temporal lag could signal a slower recruitment of the underlying neural mechanisms required for behavioral inhibition and threat evaluation^79,80^.

Another interesting aspect of our study lies in the assessment of behavioral flexibility and adaptability through frequency and transition metrics^81,82^. ELS animals showed reduced Simpson diversity and increased cumulative usage scores, indicating reliance on a narrower behavioral repertoire. At the same time, increased recurrence rates and decreased Markov entropy suggest more predictable, stereotyped behavior sequences in ELS mice. Interestingly, the LBN model is characterized by an increased entropy of maternal behaviors during the ELS exposure, suggesting that unpredictability rather than quantity, may drive the long-term effects in the offspring^83^. Therefore, the subsequent reduction in entropy in the ELS offspring may reflect a compensatory adaptation to early-life unpredictability, with these structured, repeatable action sequences possibly indicating not a just loss of flexibility but a context-specific response to early environmental uncertainty^10^. These findings in entropy are also in line with work in other stress models showing reduced behavioral complexity of stressed mice in the social interaction task^69^. While a decrease in entropy can reflect learning and improved performance in other contexts (e.g. marmosets during foraging;^84^), our data suggest that without a concurrent maintenance of behavioral variety (reflected in reduced Simpson diversity) this pattern may instead indicate rigidity.

This behavioral rigidity might indicate a loss of flexibility that impairs the animal’s capacity to adjust to new or evolving threats. Control animals display more varied and spontaneous sequences. In contrast, while transitions between behaviors still occur, ELS animals show a tendency to cycle through a narrow set of patterns. Such stereotypy is common in stress and anxiety-related conditions^85,86,87,88^, where perseveration replaces more adaptable coping mechanisms.

However, this does not necessarily mean ELS-induced behavioral rigidity is inherently maladaptive. Instead, ELS may foster coping strategies that favour stability and predictability. In environments that are stable or familiar, such predictable behavioral responses may act as a risk-aversive mechanism that reduces uncertainty and prioritizes safety. However, when faced with challenges that require flexibility (such as a subsequent stressor) rigid responses may prove limiting. In this sense, reduced behavioral entropy reflects an adaptive trade-off rather than a deficit: a shift toward predictable behavior that may work well in stable environments but less so when demands change, in line with the match/mismatch hypothesis^89,90,91,92^. In the future it will be important to investigate behavioral repertoires of control and ELS mice also in more naturalistic behavioral and stressful contexts^93,94^.

Next, we adapted the von Ziegler et al. (2024) approach, which uses dimensionality reduction on transition matrices and pairwise distance metrics to compress complex behavioral data into a single measure that they named as *behavioral flow*. Given that our FC paradigm involves repeated shocks, we focused on capturing dynamics over time rather than just transition counts. We binned the behavior into time intervals, computed pairwise distances, and used MDS to obtain a two-dimensional profile for each animal. From this space, we derived a score based on the log ratio of each animal’s Euclidean distance to the ELS median versus the control median. This streamlined metric allowed us to classify ELS animals, with those scoring closer to the control group. These mice were considered as “resilient”.

Approximately 30% of the offspring exposed to ELS was classified as resilient. This percentage matches several human reports of resilience after ELS (^95^: maintained positive peer relationships, completed high school and showed no serious mental illness by age thirty;^96^: free of psychiatric disorder, substance abuse and criminal convictions with stable employment and close relationships in early adulthood;^97^: behavioral scores at age eleven within the range of non-exposed children), but this percentage varies significantly across human cohorts^98,99,100,101^.

In our study, resilient animals not only showed higher and prolonged levels of freezing behavior but also developed behavioral profiles over time that were more comparable to control mice, including greater diversity, increased variability, unpredictability, and reduced repetitiveness in their actions, reflecting less stereotypy. Notably, this subgroup even surpassed control animals in certain behavioral aspects, such as freezing frequency and dynamics, suggesting a more adaptive or context-appropriate response to stressors. This observation supports the idea that freezing is not merely a passive fear response, but part of an active, coordinated coping strategy that can enhance adaptability^76^. Our data suggests that the enhanced freezing over time seems to play a important role in driving resilience in the current experimental context.

During FC, the “vulnerable” ELS animals exhibited highly predictable behavior, cycling through a narrow set of responses and defaulting to familiar strategies rather than exploring alternatives. Meanwhile, resilient animals, although not fully overcoming the reduction in behavioral entropy (a potential cost), compensated by enhancing their limited repertoire and exhibiting less recurrent behavioral sequences. This may reflect flexibility to rapidly adjust their behavioral responses to changing environmental demands. Increased behavioral flexibility in stressed individuals upon a second stressor has been considered a signature of resilience^102,103,104^. Interestingly, in some cases, this flexibility is linked to shifts in coping strategies from active to passive^105^.

Finally, it will be important for future studies to consider behavioral variability within the control group. While this study focuses on the resilience of ELS animals, assessing potential vulnerabilities in controls will help ensure that differences between groups reflect true effects of the intervention.

Due to the combination of different datasets in this study (with e.g. different endpoints), we were unable to find a physiological or neurobiological correlate of our behavioral signature for resilience. Although underpowered to make definitive claims, we have preliminary data suggesting that resilient animals also have immediate-early gene cell numbers closer to controls^47^. It is also suggested that interneurons and the excitation/inhibition (E/I) balance could mediate adaptation upon ELS and glucocorticoids^106^. Understanding the underlying mechanisms that allow for this resilience-whether through genetic and epigenetic predisposition, moderate neural plasticity, or other factors such as the immune system or the gut microbiome^107,103^-could provide important insights into potential interventions for individuals who have experienced early-life adversity.

Rather than viewing resilience and vulnerability as strictly dichotomous outcomes, our study suggests that these concepts can be defined as points along a continuum, shaped by multiple interacting factors, including inherent “mousenality”: the individual behavioral traits and tendencies unique to each mouse. In this view, the same experimental manipulation can lead to different outcomes depending on the subject. Therefore, efforts should be made to improve individual differences analysis, recognizing that variation across subjects can offer critical insight rather than being dismissed as noise or lost in the average per group.

Animal models offer valuable insights for studying stress-related disorders, yet significant translational challenges persist due to a reductionist approach of, among others, behavioral analysis^108^. Traditional laboratory tasks, designed to probe a specific stress-induced phenotype, may fail to capture ethologically relevant behavioral diversity. A major shift in behavioral neuroscience came with the introduction of pose estimation tools like DeepLabCut^23^, which allowed high-throughput tracking of animals. Since then, several other open-source systems have become available, including DeepPoseKit^109^, or SLEAP^110^. These advances have driven interest in computational methods, particularly machine learning, for refining behavioral analysis^108,111,112^.

Deep phenotyping methods can be broadly divided into two categories: supervised and unsupervised classification. Supervised methods rely on labeled data and either rule-based logic or machine learning models to identify specific behaviors. For instance, we have implemented SimBA^24^ to automatically score freezing behavior in the FC paradigm using a random forest classifier^46,47^. Other pipelines like SIPEC^113^, MARS^28^ and specifically for FC, BehaviorDEPOT^114^ use similar strategies, integrating tracking data with predefined-manual-behavioral annotation. While these tools are useful, they depend on pre-labeled data, limiting their flexibility for exploratory or novel behavioral insights.

Unsupervised classification methods, on the other hand, extract patterns without relying on predefined behavioral categories, which makes them powerful for hypothesis generation. B-SOiD^31^ uses nonlinear embeddings of pose trajectories to segment behavior from movement, and A-SOiD applies a similar strategy enhanced by active learning to reduce manual annotation. VAME^33^ learns latent representations of behavioral dynamics using a variational autoencoder with recurrent networks, while DeepOF^27^ and CEBRA^34^ leverage deep and contrastive learning, respectively, to cluster and differentiate behavioral motifs. These systems offer powerful ways to uncover structure in continuous behavior and have pushed the field toward more data-driven behavioral science.

Nevertheless, unsupervised methods have their limitations. These methodologies can be prone to overfitting, difficult to validate, likely to generate patterns lacking clear behavioral meaning, and unable to capture temporal dynamics^115^. To partially address this, we used keypoint MoSeq^34^ as it uses an autoregressive hidden Markov model, allowing it to capture temporal structure and segment behavior into discrete, interpretable syllables. This was especially important in our context, since FC inherently has a temporal component in which the footshocks represent experimental manipulations over the behavioral course. Keypoint MoSeq also generates a low-dimensional latent representation that preserves behavioral structure across animals and experimental sessions, facilitating consistent cross-subject comparisons. At the same time, we count with the previously reported freezing labels generated with SimBA^46,47^ as a ground truth measure to validate our unsupervised findings. In addition, we reclusterized keypoint MoSeq’s results into researcher-validated behaviors to maintain relevance and filtered out noisy clusters through visual inspection. This helped us reduce false positives and non-interpretable patterns. These clusters were further validated with explainability tools like SHAP^59^ improving transparency and interpretability as encouraged in the field^116^. Notably, some of our results overlapped with the unsupervised outcomes reported by Bordes et al. (2024), suggesting emerging reproducibility, and we propose cross-validation between datasets as a next step toward building much-needed benchmarks.

Still, overrepresentation of certain behaviors and testing too many features can inflate significance and make it hard to know which changes truly matter. Better standards for what counts as meaningful behavior are still needed. Also, when possible, predictive models should be adopted as a standard for judging the quality of behavioral representations^117^. We followed this recommendation by training a classifier to distinguish experimental groups with an overall accuracy of 62.7%, providing a quantitative readout of our behavioral models’ reliability.

Ultimately, every behavioral pipeline, whether supervised or unsupervised, must rely on choices made by the experimenter. These choices cover which behavioral features to measure, whether to analyze the behavior in the time domain or the frequency domain, what timescales to examine and whether to portray the behavior as continuous, discrete (in this case also accounting for granularity) or a mix of both^117,116^. And for that researchers should critically reflect on the ethological nature of the tests in order to enhance the validity and interpretation of their results.

## 5 Conclusion

Our study demonstrates that a previous history of ELS reconfigures the way animals behave during a fear conditioning task. Using computational pose estimation and unsupervised learning methods, we were able to capture a more detailed, time-resolved behavioral profiles that extended beyond traditional measures. We not only confirmed that ELS animals show a slower increase in freezing behavior when compared to control animals, but also we reported additional behavioral patterns showing that ELS alters the balance between passive and active coping strategies. In particular, ELS animals exhibited a narrow, rigid, repetitive, and predictable behavioral repertoire that was marked by increased durations of active behaviors such as turning and sniffing, suggesting that ELS offspring behaves more stereotypically upon a secondary challenge. Notably, while many ELS animals displayed these rigid and predictable responses, we identified a “resilient” subgroup that has comparable responses to control mice and demonstrated adaptive, flexible behavior characterized by lower entropy, greater behavioral diversity, and enhanced frequency and duration of passive behaviors compared to their vulnerable counterparts. Their adaptive trajectories, sometimes even surpassing those of control animals, may indicate a compensatory capacity that could underlie successful coping in the face of early-life adversity. These insights emphasize the need to move beyond the traditional, monolithic view of freezing in fear conditioning toward a broader analysis that encompasses the full spectrum of behaviors and metrics such as diversity and entropy, a shift that holds broader implications for ethologically based behavioral analysis and future research on stress-related disorders.

## Acknowledgements

We thank SURF for their computational support through an NWO/EINF grant [EINF-11916]. We especially thank Carlos Teijeiro Barjas for guiding us through SURF’s services. We also thank Dora Gözükara and Kieran Carrigg and Gamze Kantar (Neural Coding Lab, Donders Institute for Brain, Cognition and Behaviour, Radboud University) for their valuable discussions on the project. Finally, we thank Rodrigo García Valiente and Nora Leger for their support during the preparation of this manuscript.

## Author contribution

**Jeniffer Sanguino Gómez**: Conceptualization, Data Curation, Formal Analysis, Funding Acquisition, Investigation, Project Administration, Resources, Supervision, Validation, Visualization, Writing - Original Draft, Writing - Review & Editing. **Umut Güçlü**: Conceptualization, Methodology, Resources, Supervision, Writing - Review & Editing. **Harm J. Krugers**: Funding Acquisition, Resources, Supervision, Writing - Review & Editing. **Antonio Lozano**: Conceptualization, Data Curation, Formal Analysis, Funding Acquisition, Methodology, Project Administration, Resources, Software, Supervision, Validation, Visualization, Writing - Original Draft, Writing - Review & Editing. All authors have read and agreed to the published version of the manuscript.

