## Supplementary_material for "Coping strategies dynamics and resilience profiles after early life stress revealed by behavioral sequencing"

### Appendix

| Behavior | Definition |
|---|---|
| Freezing | Absence of all voluntary body movement except for respiration. No displacement of the center of mass; posture held in fixed position. Common in response to perceived threat or during heightened vigilance. |
| Sniffing | Repetitive, rapid head and snout movements directed toward a stimulus. Minimal forward movement of the body. Typically occurs during investigation or environmental exploration. |
| Grooming | Repetitive, patterned movements involving forepaws or head contacting the face or body. Includes licking, rubbing, or scratching actions. Often observed during self-maintenance, recovery from arousal, or transitions between behavioral states. |
| Turn | Rotation of the body's orientation through paw repositioning. May involve pivoting or stepping. Center of mass rotates in place or shifts laterally. Common during orientation, exploration, or avoidance. |
| Locomotion | Coordinated, alternating limb movements that result in forward or lateral displacement of the body's center of mass. Used in exploration, approach, or escape behaviors. |
| Climbing | Vertical or diagonal movement along a surface, using forelimbs and hindlimbs in alternation to elevate the body. Often involves elongation and gripping. Seen during exploration, escape attempts, or interaction with vertical structures. |
| Jump | Rapid, forceful extension of all four limbs causing the body to lift off the ground. Center of mass becomes airborne with no limb contact. Typically occurs during escape, gap crossing, or rapid navigation of terrain. |

**Supplementary Table 1**. Definitions combining physical structure and ethological context of the seven ethologically relevant behaviors derived from keypoint MoSeq syllable clustering, which constitute the categories used for analysis in this study.

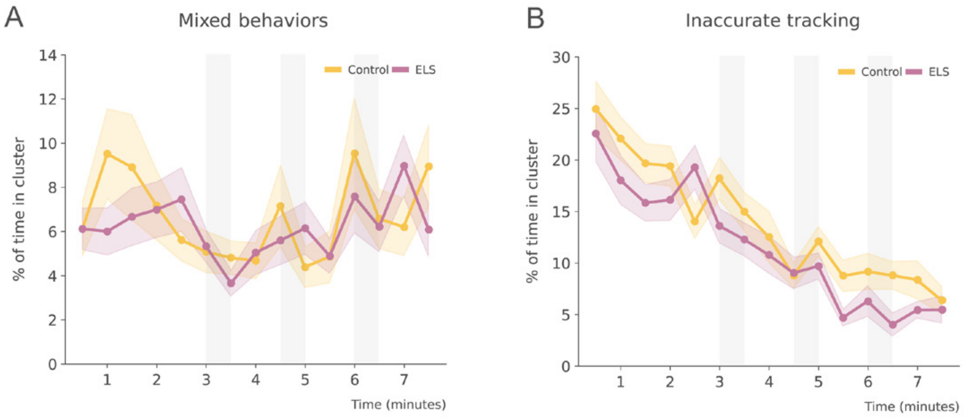


**Supplementary Figure 1**. ELS does not alter non-ethological cluster usage during fear conditioning. **A**. Lineplots showing the "mixed behaviors" cluster (mean % ± SEM) over time during FC in control and ELS animals. The percentage of mixed behaviors is equal between groups. **B**. Lineplots showing the "inaccurate tracking" cluster (mean % ± SEM) over time during FC in control and ELS animals. Inaccurate tracking was unaffected by condition. Shaded areas represent tone and footshock epochs. $N_{Control} = 41$, $N_{ELS} = 41$.

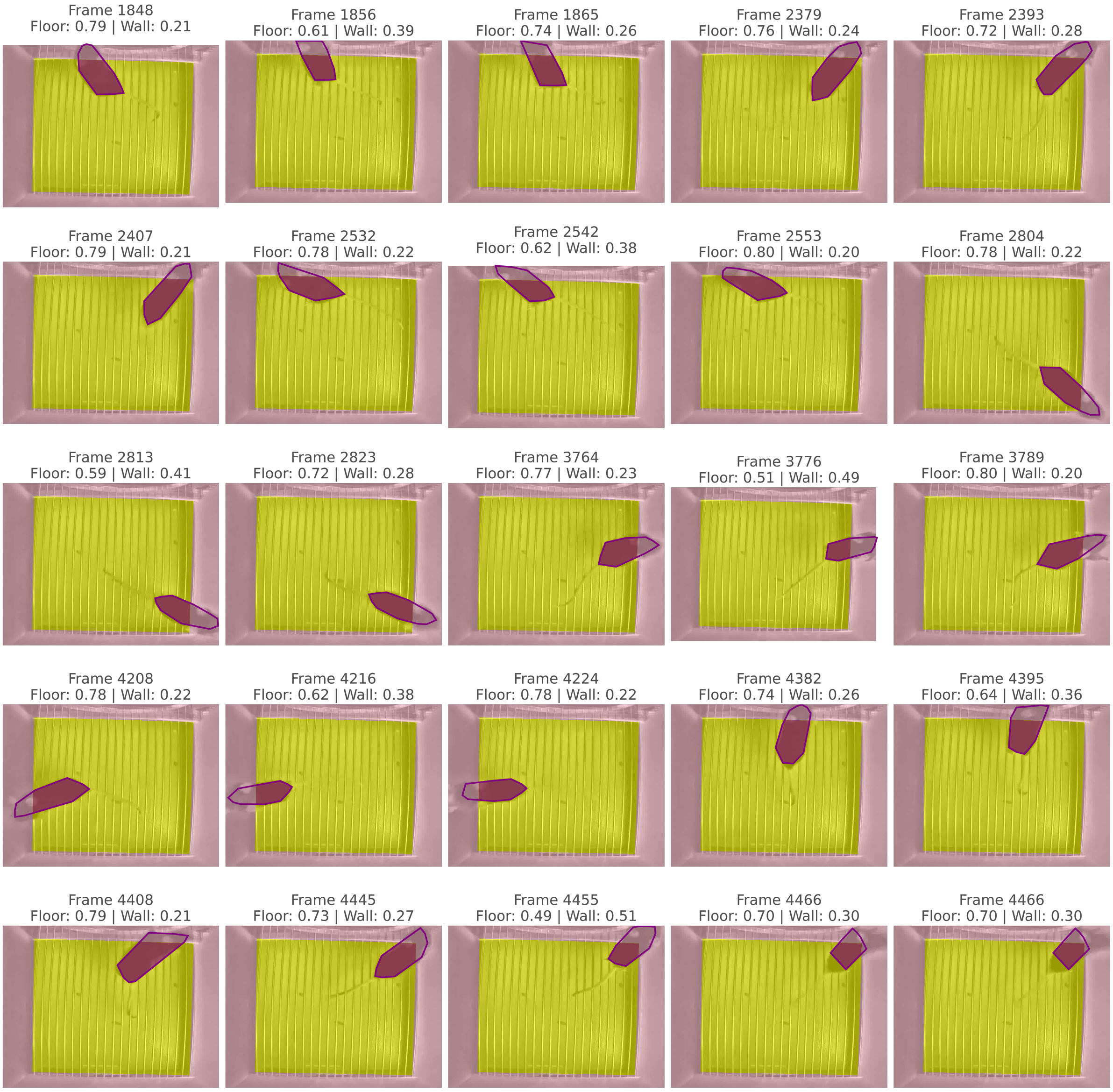


**Supplementary Figure 2**. Convex-hull floor-overlap metric allows for climbing identification missed by keypoint MoSeq. Twenty-five representative top-down frames showing how climbing is resolved from two-dimensional video. A binary mask derived from manually annotated cage corners partitions the arena into floor (light yellow) and walls (mauve). The convex hull of the animal's DeepLabCut keypoints is overlaid in dark purple. Numbers above each frame report the proportion of hull area on the floor and on the walls. As the mouse ascends the wall the hull shifts upward, floor overlap falls below 0.8, and wall overlap rises, flagging a climbing episode that keypoint MoSeq alone could not label reliably.

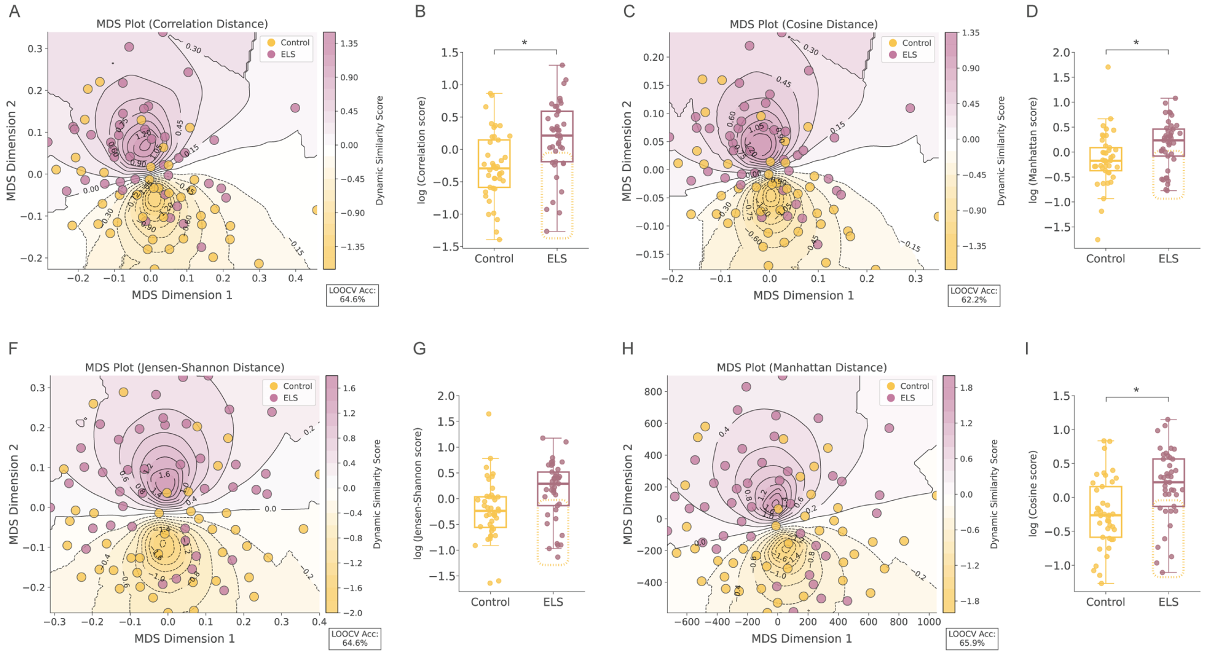


**Supplementary Figure 3**. Alternative distance metrics consistently separate ELS and control mice in a behavioral-dynamics space. **A**. MDS plot of the correlation-distance matrix computed from pairwise comparisons of each animal's behavioral-dynamics profiles. Each dot is the two-dimensional embedding of one mouse, positioned so that inter-dot distances reflect the original correlation distances in feature space. Shaded ellipses represent each group's median (dark core = median, fading edges = dispersion). Leave-one-out cross-validation accuracy (LOOCV Acc) is indicated in the bottom corner. **B**. Boxplots of the log-transformed correlation-distance scores (mean ± SEM) for control versus ELS animals. ELS mice differ significantly from controls. Square markers enclose ELS animals with scores below zero, classified as resilient. **C**. MDS plot of the Cosine-distance matrix computed from the same behavioral-dynamics profiles. Each dot is the two-dimensional embedding of one mouse, positioned so that inter-dot distances reflect the original cosine distances in feature space. Shaded ellipses represent each group's median (dark core = median, fading edges = dispersion). Leave-one-out cross-validation accuracy (LOOCV Acc) is indicated in the bottom corner. **D**. Boxplots of the log-transformed Cosine-distance scores (mean ± SEM) for control versus ELS animals. ELS mice differ significantly from controls. Square markers enclose resilient ELS animals. **E**. MDS plot of the Jensen–Shannon-distance matrix computed from the behavioral-dynamics profiles. Each dot is the two-dimensional embedding of one mouse, positioned so that inter-dot distances reflect the original Jensen–Shannon distances in feature space. Shaded ellipses represent each group's median (dark core = median, fading edges = dispersion). Leave-one-out cross-validation accuracy (LOOCV Acc) is indicated in the bottom corner. **F**. Boxplots of the log-transformed Jensen–Shannon-distance scores (mean ± SEM) for control versus ELS animals. No significant differences were observed between groups. **G**. MDS plot of the Manhattan-distance matrix computed from the behavioral-dynamics profiles. Each dot is the two-dimensional embedding of one mouse, positioned so that inter-dot distances reflect the original Manhattan distances in feature space. Shaded ellipses represent each group's median (dark core = median, fading edges = dispersion). Leave-one-out cross-validation accuracy (LOOCV Acc) is indicated in the bottom corner. **H**. Boxplots of the log-transformed Manhattan-distance scores (mean ± SEM) for control versus ELS animals. ELS mice differ significantly from controls. Square markers enclose resilient ELS animals. $N_{Control} = 41$, $N_{ELS} = 29$, $N_{ELS\ resilient} = 12$. * ELS effect. Effect $p \leq 0.05$.
